# Root exudates enhance the ecological fitness of a non-native PGPR and reshape rhizosphere microbiome assembly

**DOI:** 10.64898/2026.10.13.758781

**Authors:** Sangita Mondal, Dibakar Roy, Dhruba Bhattacharya, Sriparna Mondal, Utpal Bakshi, Kunal Kumar Saha, Sudipto Saha, Anupama Ghosh, Abhrajyoti Ghosh

## Abstract

Root exudates play critical roles in rhizosphere microbiome assembly. Nevertheless, the molecular mechanisms by which plant growth-promoting rhizobacteria (PGPR) perceive these signals, adapt to the rhizosphere, and influence native microbial communities remain poorly understood. Here, using maize (*Zea mays* cv. Early Golden Bantam) and the rhizobacterium *Priestia aryabhattai* AB211, we took a multi-omics approach combining metabolomics, bacterial transcriptomics and proteomics, and rhizosphere metagenomics to elucidate the mechanism underlying this process. Maize root exudates were enriched in diverse carbon and nitrogen metabolites that promoted bacterial growth and induced extensive molecular reprogramming of pathways involved in nutrient acquisition, chemotaxis, motility, stress adaptation, and plant growth promotion, thereby enhancing rhizosphere competence. In field-conditioned soil, AB211 persisted as a core member of the maize rhizosphere microbiome, promoted early seedling growth, and reshaped microbial community composition and function. Root exudates-primed (RE-primed) AB211 further enhanced the enrichment of microbial functions associated with nutrient cycling, quorum sensing, plant growth promotion, and rhizosphere fitness, highlighting the role of root exudates in modulating the rhizosphere-adaptative potential of beneficial bacteria. These findings establish a mechanistic framework linking root exudates composition with PGPR adaptation and microbiome assembly, providing a foundation for engineering ecologically adapted microbial inoculants to improve crop productivity and sustainable agriculture.

## Introduction

The rhizosphere represents the most dynamic and functionally significant interface between plant roots and soil. The microbial community of the rhizosphere is involved in diverse metabolic activities, including plant nutrition and productivity, disease suppression, and stress resilience, thereby substantially contributing to plant fitness and ecosystem functioning^1^. Despite this immense taxonomic diversity, only a subset of microbes successfully colonizes the rhizosphere, suggesting that microbiome assembly is actively governed by strong ecological selection. Understanding the intricate mechanisms underlying the recruitment and persistence of beneficial microbial communities in the rhizosphere remains a fundamental challenge in plant-microbe research.

Plant roots release a diverse array of compounds as root exudates (RE), allocating more than 21% of their photosynthetic matter in the rhizosphere^2^. RE comprises low and high-molecular-weight compounds, including amino acids, sugars, secondary metabolites, phenolics, flavonoids, organic acids, proteins, and complex carbohydrates. Hence, the rhizosphere constitutes a discrete, nutrient-rich microenvironment with physicochemical properties markedly different from those of the bulk soil^3^. Beyond serving as a nutrient resource, the rhizosphere functions as an ecological filter that selectively enriches, suppresses, or modulates the microbial community^4^. Consequently, root exudation is widely regarded as the principal driver of the ‘rhizosphere effect’^5, 6^. Primary metabolites, such as carbohydrates, amino acids, organic acids, and fatty acids, are typically released in relatively high concentrations, serving as readily available nutrients and energy sources for soil-dwelling microbes. In contrast, secondary metabolites, e.g., flavonoids, phytohormones, enzymes, sterols, and phytoalexins, are secreted at lower concentrations and strongly regulate microbial behavior and signaling, thereby promoting beneficial community assembly.

Root exudates composition is highly dynamic and is collectively influenced by specific plant age, genotype, and biotic and abiotic stress factors^7^. Variation in exudation patterns alters the physicochemical landscape of the rhizosphere and restructures the homeostasis of the rhizosphere microbiome by modulating selection pressure on resident microbial communities. For example, roots release compounds such as flavonoids, ACC, volatile organic compounds (VOCs), chemoattractants, signaling molecules, and bioactive chemicals such as inositol, cucurbitacin, and bithienyl and its derivatives. These compounds mediate the selective recruitment of beneficial microbes, suppress pathogenic activity, combat stress, and regulate microbial behavior through reciprocal communication via chemotaxis, quorum sensing, and metabolic reprogramming^8, 9^. Such observations suggest that root exudates are not simply metabolic by-products but rather represent an active communication interface for successful microbial assembly in the rhizosphere.

Among rhizosphere-inhabiting microbes, plant growth-promoting bacteria (PGPR) constitute a functionally important group capable of positively influencing plant performance by improving nutrient acquisition and phytohormone production, mitigating stress by modulating the plant stress response via induced systemic resistance (ISR), and biocontrol of detrimental phytopathogens^10, 11, 12^. Numerous studies have demonstrated that root exudates selectively stimulate PGPR lifestyle and colonization, suggesting that the successful establishment of PGPR in the rhizosphere depends on microbes’ ability to perceive and respond to plant-derived chemical signals^13^. However, the underlying mechanism of these adaptive strategies remains elusive. Few reports examine PGPR responses during beneficial plant-microbe interactions, especially with non-model PGPR strains. Transcriptomics-based studies on *Bacillus amyloliquefaciens* FZB42, *B. amyloliquefaciens* SQR9, *Burkholderia pyrrocinia* P10, suggested that pathways related to the nutrient utilization, biofilm formation, flagellar assembly, quorum sensing, and plant growth-promotion were significantly enhanced upon root exudates treatment^14, 15, 16^. These findings demonstrated that root exudates serve as environmental cues that reprogram PGPR physiology to facilitate rhizosphere adaptation and host associations.

Although transcriptomic studies have provided valuable insights into gene expression changes in PGPR upon exposure to root exudates, our understanding of how these transcriptional changes are translated into functional protein expression and cellular physiological responses remains limited. The integration of transcriptomic and proteomic analyses bridges the gap between identifying regulatory responses and their respective physiological implementations. Such systems-level interdisciplinary approaches will provide comprehensive insights into the adaptive strategies employed by PGPRs during environmental transitions and host-associated lifestyles^14, 17^. Nevertheless, multi-omics investigations of PGPR responses to plant-derived chemical signals remain scarce, especially for non-model beneficial microorganisms.

Beyond individual plant-microbe interactions, successful PGPR establishment depends on interactions with resident microbial communities^18^. Most studies on plant growth promotion have been conducted under sterile conditions, but examples of reintroducing lab-grown bacterial inocula into the field are scarce. In nature, the rhizosphere is inhabited by a complex indigenous microbial community that often resists the establishment of introduced microbial populations. Recently, researchers have focused on the effect of PGPR invasions into the native rhizo-microbiome community^9, 19^. These studies indicate that highly diverse native microbiome communities resist invasion by efficiently exploiting available resources, thereby maintaining a resource-limited environment that restricts the establishment of invading species^20^. In this competitive environment, gram-positive bacteria, especially spore-forming PGPB such as *Bacillus*, benefit from their dormant nature, which helps them evade competition from the resident microbial community. Studies revealed that the native core microbiome often possesses a “home-field advantage”, enabling it to resist colonization by the otherwise non-native beneficial microorganisms^21, 22^. Consequently, the ability of an introduced PGPR to survive in the native community depends not only on its plant-growth-promoting traits but also on its adaptive resilience, its integration into the microbial community network, and its influence on community functions^23^. Additionally, non-native PGPR, although potent under sterile conditions, lose their efficacy in natural environments^24^. These studies collectively suggest that successful rhizosphere colonization of a potent PGPR follows a two-step process. First, the root exudates modulate the physiology and behavior of beneficial microbes. Second, the adapted beneficial populations may subsequently alter microbial community composition, facilitating functional enrichment within the rhizosphere. However, the mechanistic links connecting this root exudates dynamics, bacterial adaptation, and microbiome assembly remain poorly understood.

This study aims to address this knowledge gap by investigating the molecular and ecological mechanisms underlying beneficial plant–microbe interactions during early rhizosphere colonization. Here, we have selected maize (*Zea* mays) plants as a model system and investigated the beneficial plant-microbe crosstalk mechanism using a non-model tea rhizosphere isolate, *Priestia aryabhattai* AB211. Primarily, we investigated the exudation pattern and composition of root exudates from maize (early golden bantam) across different seedling stages. Subsequently, using transcriptomic and proteomic approaches, we elucidated the mechanistic basis of root exudates-mediated adaptation in *Priestia aryabhattai* AB211, which facilitates successful root colonization in the early stages of the beneficial interaction. Finally, using a native maize rhizosphere soil system, we evaluated the impact of AB211 and a root-exudates-primed (RE-primed) AB211 amendment on plant growth, microbial community dynamics, and functional enrichment of the beneficial microbial population within the rhizosphere. By integrating multi-omics analyses with microbiome ecology, this study highlights a system-level framework linking rhizosphere effects to bacterial adaptation and beneficial microbiome assembly, thereby elucidating the ecological and agricultural significance of PGPR application in agriculturally valuable crop systems.

## Results

### Maize root exudates composition

Maize root exudates were collected in the aqueous phase and analyzed by HPLC to quantify monosaccharides, oligosaccharides, amino acids, and organic acids (Fig. 1, Supplementary Table S1). Among diverse metabolites, glucose and melibiose were found to be abundant mono– and oligosaccharides, followed by an adequate amount of isomaltose, maltose, fructose, lactose and others (Fig. 1B). Among the detected organic acids, malonic acid and lactic acid were present at the highest concentrations (Fig. 1C). Trace level of amino acids such as alanine, aspartic acid, leucine, glutamic acid, and others were also detected in the root exudates (Fig. 1D).

**Fig. 1.**
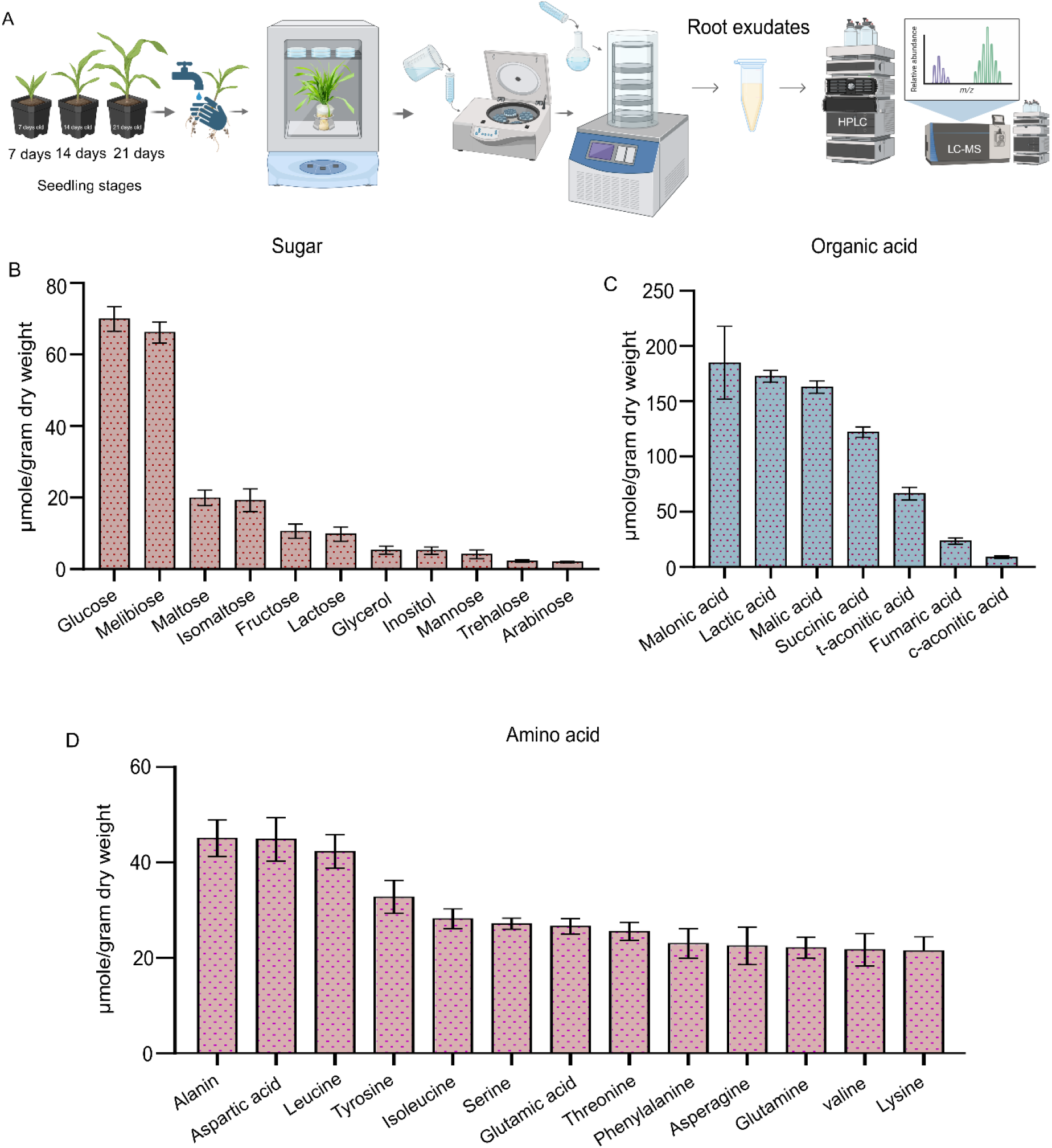
Schematic representation of root exudates extraction from three different seedling stages (07, 14, and 21 days old) of maize. Estimation of primary nutrient sources like sugars (A), organic acids (B), and amino acids (C) in root exudates through HPLC. Error bars represent the standard error of mean (n=3 replicates).

To investigate the comparative metabolome profile across three seedling stages, untargeted LC-MS/MS was performed. The raw metabolome data were processed and identified using the ChemSpider and PlantCyc databases^25^. A total of 1586 m/z features were obtained from the untargeted detection. Based on their abundances, highly abundant metabolites across three consecutive days were manually screened and presented in Supplementary Fig. 1A. Most of the abundant compounds include plant hormones, such as methyl jasmonate and trans-zeatin, long-chain fatty acids (9, 10-, 12– 13– diepoxyoctadecanoate), and antimicrobial compounds (coumarin and associated derivatives). The stage-specific variation in their abundance at 7, 14, and 21 days suggests that plants diligently modulate the composition of root exudates throughout development to meet physiological and ecological requirements.

### Planktonic growth pattern of AB211 upon exposure to root exudates

The significant influence of crude root exudates on the growth behavior of AB211 was evident from the growth curve analysis (Supplementary Fig. 1B). The root exudates treatment significantly influenced the growth rate of AB211 compared with the untreated control. Interestingly, the growth behavior followed a similar trend across different root exudates concentrations, viz. 0.25 mg/ml and 0.5 mg/ml with no significant alterations. Similarly, chemotaxis and motility behavior were profoundly influenced by exposure to root exudates, especially during the exponential phase (O.D._600_ = 1.0) (Supplementary Fig. 2A&B). Notably, biofilm formation was significantly increased upon RE exposure, and enhanced production was observed during the stationary phase (O.D._600_= 2.0) rather than the exponential phase (Supplementary Fig. 2C).

Plant growth-promotion attributes, viz., indole-3-acetic acid (IAA) and siderophores, were also enhanced in AB211 across both growth phases in the presence of root exudates. IAA production increased by more than 2-fold both during the exponential and the stationary phase following RE treatment (Supplementary Fig. 2D). Similarly, quantification of total siderophore production by observing the iron chelation increased significantly from 48.36% to 63.38% during the exponential phase (Supplementary Fig. 2E). Additionally, siderophore typing of AB211 revealed the possible presence of both hydroxamate and carboxylate type siderophores in culture. However, no catecholate-type siderophore was detected. Quantitative assessment revealed hydroxamate-type siderophore production increased from 9.74 to 10.94 µg mL⁻¹ and 10.39 to 11.32 µg mL⁻¹ during the exponential and stationary phases, respectively, under RE treatment (Supplementary Fig. 2F). Similarly, a significant increase in carboxylate-type siderophore production was observed upon RE treatment both during exponential (from 35.80 to 95.89 µg mL⁻¹) and stationary phase (from 51.87 to 108.5 µg mL⁻¹) (Supplementary Fig.2G).

**Fig. 2.**
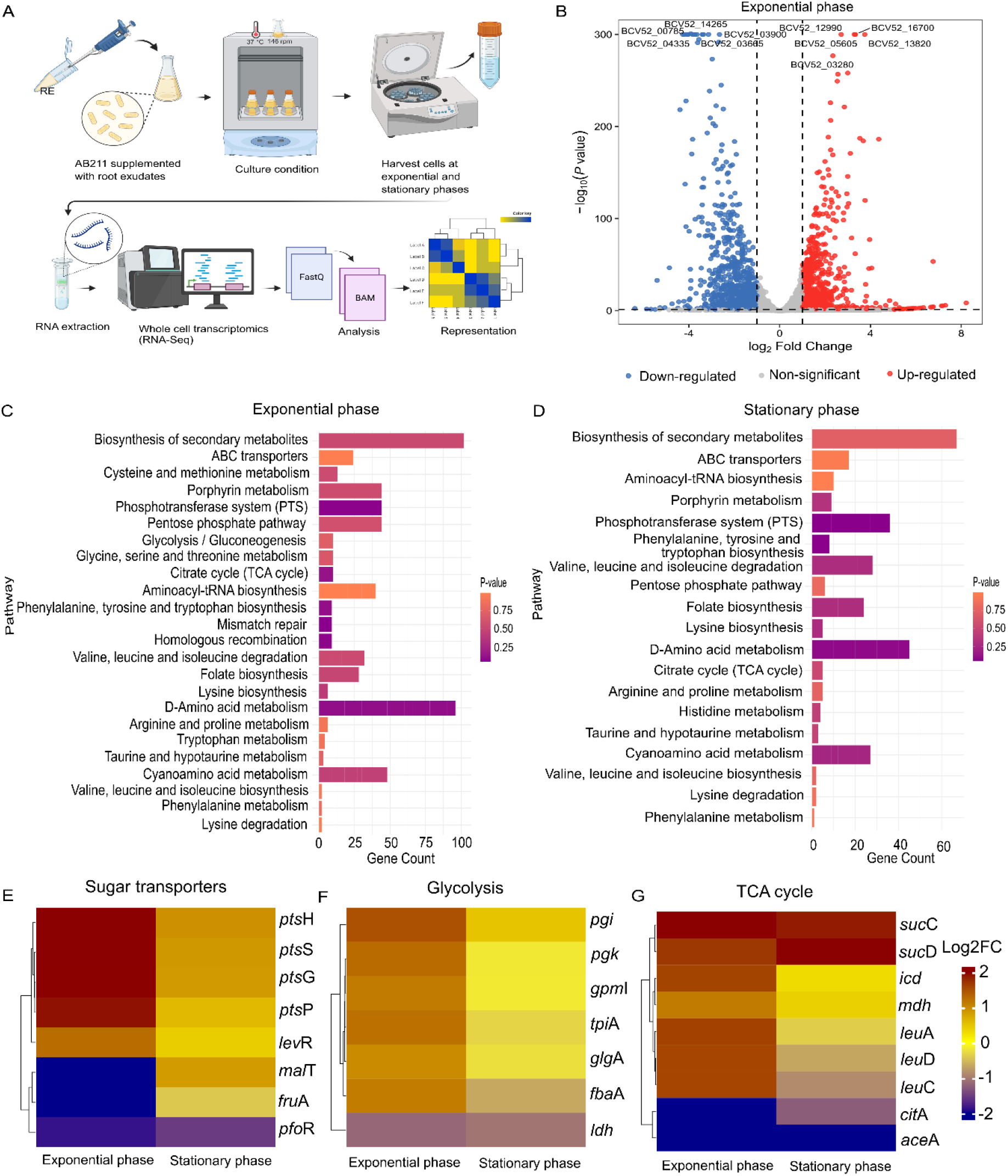
Schematic representation of transcriptomics sample preparation during exponential and stationary growth phases of *Priestia aryabhattai* AB211 upon root exudates (0.25 mg/ml) treatment. (A). The volcano plot represents the total number of differentially expressed genes during the exponential phase (B). The KEGG pathway enrichment of significant DEGs from exponential (C) and stationary (D) phases. The heatmap shows the expression patterns of significantly altered genes involved in sugar transporters (E), glycolysis (F), and TCA cycle (G). The log_2_ fold change values are represented by the provided color code (log_2_ fold change ≥ ±1, pvalue ≤0.05).

### Root exudates induce extensive transcriptional reprogramming in AB211

To understand how root exudates reshape the physiology of AB211, RNA-sequencing was performed during exponential and stationary growth phases following supplementation with 0.25 mg/ml root exudates (Fig. 2A). Transcriptome sequencing results revealed that a total of 361 genes (log2 fold change ± 1, p-value ≤ 0.05) showed significant alteration during both exponential and stationary phases (Supplementary Table S2). However, root exudates elicited a markedly stronger transcriptional response during the exponential phase than during the stationary growth phase. Overall, 1627 genes were differentially expressed in AB211 upon root exudates treatment (log2 fold change ± 1, p-value ≤ 0.05) during the exponential phase, while 781 genes showed significant alteration (log2 fold change ± 1, p-value ≤ 0.05) at the stationary phase. This gene expression pattern suggests that the exponential phase captures the primary impact of root exudates on the molecular machinery of AB211. Hence, we focused on the significantly differentially expressed genes (DEGs) identified during the exponential phase to investigate the underlying molecular mechanisms in response to exposure to root exudates (Fig. 2B, Supplementary Table S3).

The functional Clusters of Orthologous Group (COG) category of significantly up-regulated genes revealed that 16% genes were involved in amino acid metabolism, followed by genes whose products are involved in inorganic ion metabolism (8.4%), transcription (7.2%), coenzyme metabolism (5.4%), energy production (5.4%) and translation (5.4%) (Supplementary Fig. 3A & B). Among the up-regulated genes in the exponential phase, a total of 740 genes could be mapped using Eggnogmapper, including 163 with unknown functions. Around 544 mapped genes were categorized into various cellular functions, including amino acid metabolism, and translation, energy metabolism, signal transduction, intracellular trafficking (Supplementary Fig. 3B). KEGG pathway enrichment analysis revealed that pathways related to amino acid biosynthesis and metabolism, the phosphotransferase system (PTS), and the TCA cycle were consistently enriched in both the exponential and stationary phases (Fig. 2C and D). Collectively, these results indicate that root exudates predominantly influence actively growing bacterial cells and induce extensive metabolic reprogramming during the initial stages of rhizosphere colonization.

### Root exudates reprogramme nutrient uptake and central metabolism of AB211

Root exudates are enriched with diverse carbon and nitrogen metabolites, serving as a primary nutrient source for rhizosphere microbes. Sugar phosphate transporters are vital machinery for nutrient uptake and utilization. Extensive remodeling of the phosphoenolpyruvate-dependent sugar phosphotransferase system (PTS) was observed (Fig. 2E). Significant upregulation of PTS operon genes (including *pts*H (BCV52_22395): 2.8-fold, *pts*S (BCV52_09115): 2.28-fold, and *pts*G (BCV52_22400): 1.9-fold), together with significant alterations in substrate-specific PTS permeases, such as down-regulation of fructose transporters (*fru*A, BCV52_18165: –2.7-fold), indicated selective uptake and transport of carbohydrates (Fig. 2E).

Among the carbohydrate metabolism pathways, 13 genes showed significant upregulation (p-value < 0.05) during the exponential phase (Fig. 2F and G). Notably, upregulation of the majority of genes covering the entire glycolysis and TCA cycle, including *mdh* (malate dehydrogenase, BCV52_05400), *pgi* (glucose-6-phosphate isomerase, BCV52_02945), *gpm*I (phosphoglycerate mutase, BCV52_03510), *tpi*A (triose-phosphate isomerase, BCV52_03515), and *pgk* (phosphoglycerate kinase, BCV52_03520), suggests an enhanced nutrient utilization capacity of AB211 in the presence of root exudates. These results corroborate the role of root exudates as nutrient-rich substrates that support the metabolic adaptation of rhizobacteria, thereby promoting successful plant-rhizobacterial interactions. Together, these findings indicate that root exudates function as a metabolically favored carbon source that reprograms the central carbon metabolism of PGPR.

### Root exudates activate chemotactic movement and motility of AB211 towards the rhizosphere

The first step in plant-PGPR interaction involves sensing signals from root exudates, a process known as chemotaxis. The bacterial chemotaxis response pathway is initiated by the reception of chemical signals via methyl-accepting chemotaxis protein (MCP) receptors, leading to upregulation of downstream genes involved in bacterial chemotaxis^26^. In *Bacillus*, the signal perceived by MCPs is transferred to the CheA protein to activate its kinase function, followed by rapid phosphorylation of CheY of a two-component system. This phosphorylated CheY interacts with the flagellar motor protein to change the direction of bacterial movement ^27^. The transcriptome data revealed significant expression of genes encoding methyl-accepting chemotaxis protein (*mcp*C, BCV52_01900) and kinase group of phosphorylating protein (*che*A, BCV52_08355) during the exponential phase (Fig. 3A, Supplementary Fig. 4A), including coordinated upregulation of other chemotaxis signaling pathway genes (*che*D, BCV52_08365: 1.6-fold, *che*W, BCV52_08360: 2.3-fold, and *che*R, BCV52_07565: 1.2-fold) (Supplementary Table S3, Supplementary Fig.4A).

**Fig. 3.**
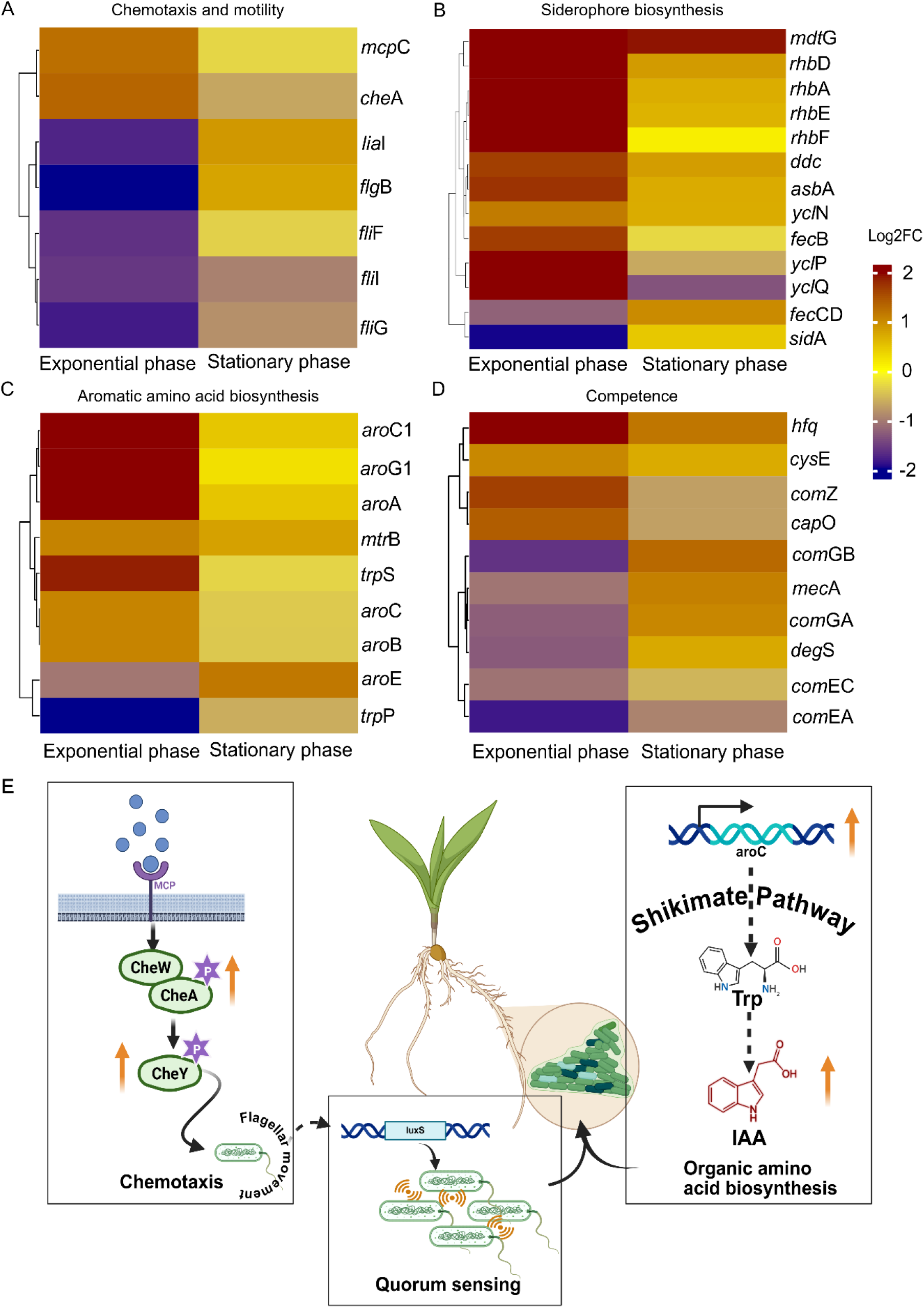
Heatmap showing the expression patterns of significantly altered genes involved in chemotaxis. (A), siderophore biosynthesis (B), aromatic amino acid biosynthesis (C), and competence-related functions (D). The log_2_ fold change values are represented by the provided color code (log_2_ fold change ≥ ±1, pvalue ≤0.05). (E) depicts the probable model of root exudate-mediated physiological reprogramming of AB211, including chemo-sensing of root exudate signals following activation of regulatory genes involved in chemotaxis and motility, thereby influencing quorum sensing and phyto-beneficial functions.

**Fig. 4.**
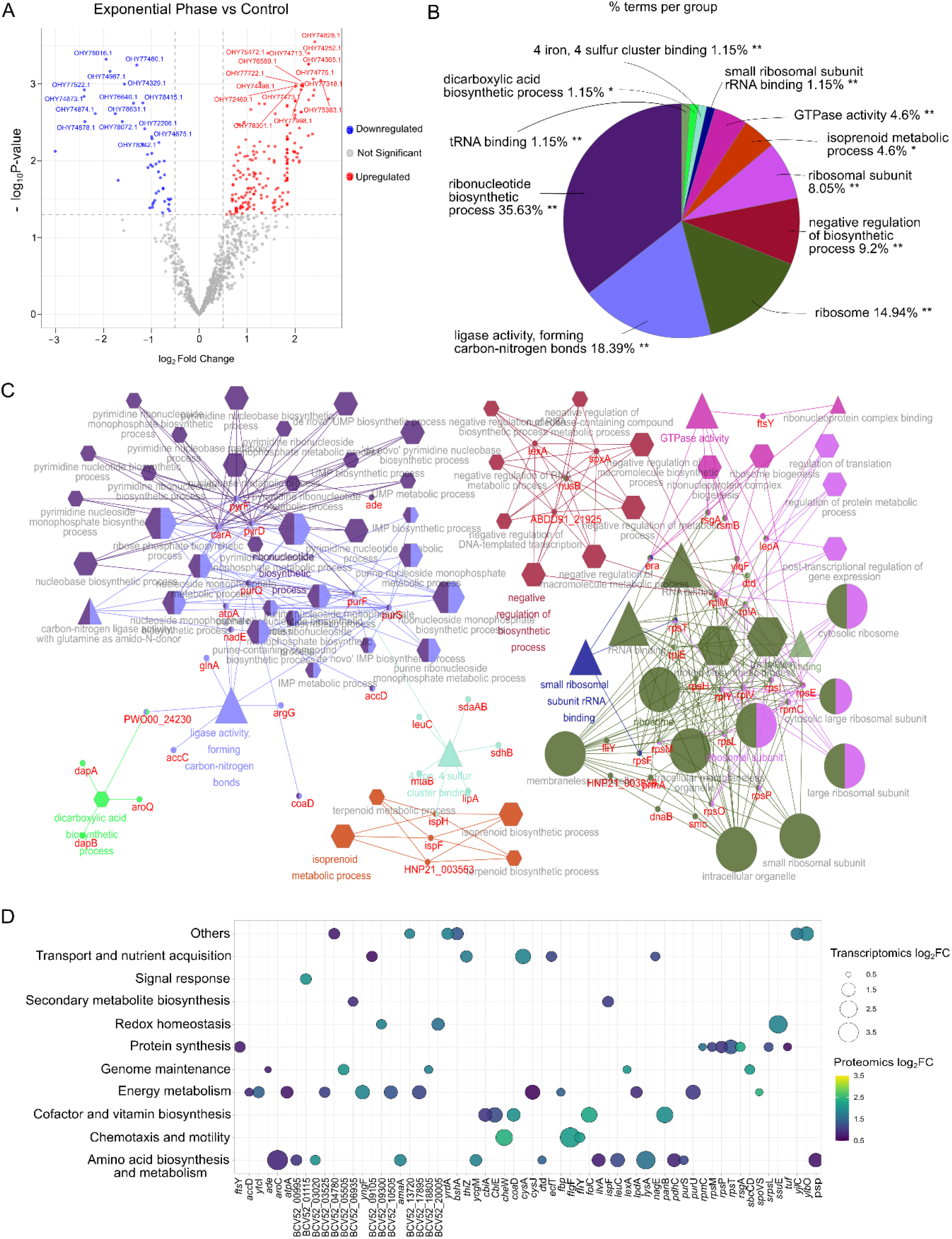
Alteration in the proteome profile during the exponential phase of AB211 (Log fold change ≥ 0.5, p-value ≤0.05). The volcano plot shows the differential protein expression profiles in AB211 in response to root exudate. (A). The pie chart depicts the upregulated pathways in the exponential phase, as identified by GO analysis (B). The interaction network shows the upregulated genes and pathways, identified through functional enrichment analysis using ClueGo, a Cytoscape plugin. Red nodes indicate upregulated genes. The same color coding was applied to both pathway nodes and their connecting edges to emphasize crosstalk between pathways and genes, using the Benjamini-Hochberg correction. Pathway enrichment analysis and visualization were performed in Cytoscape (version 3.10.2). Node shapes (triangle, circle, and hexagon) indicate the different GO ontologies: Molecular Function, Cellular Component, and Biological Process (C). Bubble plot showing concordant expression pattern of biologically significant proteins (Log fold change ≥ 0.5, p-value ≤0.05) and their encoding genes upregulated in proteome and transcriptome (Log fold change ≥ 0.5, p-value ≤0.05) (D).

Flagellar function-associated genes initiate the concordant activation of bacterial motility. Multiple homologs of genes regulating flagellar biosynthesis and functions revealed growth phase-dependent interesting expression patterns (Fig. 3A). Flagellar assembly (*fli*W, BCV52_03820 and *flg*M, BCV52_03840), (*mot*B, BCV52_18680) encoding flagellar motor protein, and basal rod protein (*flg*G, BCV52_08280) were extensively up-regulated only during the exponential phase, following non-significant expression during the stationary phase (Supplementary Table S4, Supplementary Fig. 4A). Significant (p-value < 0.05) expression of biologically relevant homologs of the *mot*B gene suggested chemotactic movement of AB211 towards the rhizosphere environment (Supplementary Fig. 4A).

Real-time PCR also showed a similar expression pattern for these genes, including *che*A, *che*W, *mcp*C, *mot*B, *fli*Y, and *flg*G (Supplementary Fig. 4A), which strongly indicated enhanced environmental sensing accompanied by reorganization of the motility apparatus. These findings confirm that root exudates stimulate directional movement by selectively activating chemotactic signaling and the motility machineries, rather than by a generalized stimulation of flagellar biogenesis.

### Root exudates enhance plant growth-promoting functions

AB211 is a potent PGPR, as revealed by its *in vitro* and *in vivo* phyto-beneficial activities. Its genome also encodes numerous PGP traits, such as phosphate solubilization, phytohormone production, biofilm formation, and siderophore production^28^. The genomic and transcriptional profiles of AB211 further supported the physiological responses linking plant growth-promotion traits. The genome contained a diverse group of siderophore biosynthesis and ferric-siderophore complex transport genes. Transcriptomic analysis, coupled with in-depth gene mining, demonstrated that root exudates significantly upregulated the expression of all the above-mentioned siderophore-associated genes (Fig. 3B) in both the exponential and stationary phases. These include genes encoding diaminobutyrate–2– oxoglutarate aminotransferase (*rhbA,* BCV52_08865) and L-2,4-diaminobutyrate decarboxylase (*ddc,* BCV52_08870) involved in biosynthesis of 1,3-diaminopropane, which serves as a primary precursor for carboxylate or mixed hydroxamate-carboxylate type siderophores like rhizobactin or schizokinen (Fig. 3B)^29, 30, 31^. Additionally, the AB211 genome also contains genes encoding siderophore biosynthesis protein (*asbA,* BCV52_08875), lysine 6-monooxygenase (*rhbE.* BCV52_08885), Lysine N-acyltransferase (*rhbD,* BCV52_08880), and ornithine monooxygenase (*sidA,* BCV52_02410) involved in biosynthesis of mixed carboxylate-hydroxamate siderophores like aerobactin and rhizobactin (Fig. 3B). Moreover, AB211 genome also contain several genes encoding ferric-siderophore complex transport proteins including, components of ferric-citrate siderophore uptake ABC transporter system (*fecCD*, BCV52_03330; *fecB*, BCV52_03335) and ferric-petrobactin siderophore uptake ABC transporter (*ycl*N_BCV52_12975, *ycl*O_BCV52_12980, *ycl*P_BCV52_12985 and *ycl*Q_BCV52_12990) and possible siderophore exporter MFS transporter (*mdt*G_BCV52_08890) (Fig. 3B)^29, 31, 32, 33^. Nonetheless, qRT-PCR further supported the elevated expression of the siderophore biosynthesis protein-encoding gene *rhb*F (BCV52_08895) and the ferric-siderophore complex-binding protein-coding gene *fec*B (BCV52_03335) upon root exudates treatment of AB211 growing in the exponential phase (Supplementary Fig 4B).

Additionally, PGPR produce IAA, a well-characterized phytohormone essential for plant development. AB211 was previously screened for IAA production and found to typically produce IAA with increasing concentrations of the precursor molecule tryptophan^28^. The application of root exudates enhanced the expression of aromatic amino acid biosynthesis pathway-related genes, which play a critical role in IAA production. Notably, the essential genes that catalyze the formation of 3-deoxy-D-aribino-hept-2-ulosonate-7-phosphate (*aro*G1, BCV52_13820), chorismate synthase (*aro*C1, BCV52_13825), 3– phosphoshikimate 1-carboxyvinyltransferase (*aro*A, BCV52_13835) exhibited more than 2-fold induced expression on Log_2_ scale (Fig. 3C). This indicated enhanced synthesis of chorismate, the common precursor of aromatic amino acids and numerous secondary metabolites. Chorismate synthase (*aro*C, BCV52_07570) converts 5-enolpyruvylshikimate-3-phosphate to chorismate. This initial branch-point metabolite increases the biosynthesis of aromatic amino acids required for IAA biosynthesis^34, 35^. The isochorismatase gene is a necessary component of the chorismate-dependent pathway and is also involved in siderophore production^36^. Apart from siderophore production, isochorismatase also influences the availability of downstream precursors, such as tryptophan, thereby increasing the IAA biosynthetic capacity of PGPR. Likewise, upregulation of nitrilase, which catalyzes the conversion of indole-3-acetonitrile via the nitrile-dependent auxin biosynthesis pathway, suggests IAA production in AB211. The expression patterns of *aro*C (BCV52_07570) and isochorismatase (BCV52_06230) have been validated by real-time PCR (Supplementary Fig. 4C). Thus, the RE-induced enhancement of IAA and siderophore production was accompanied by the expression of genes associated with siderophore biosynthesis and iron uptake, demonstrating coordinated plant-responsive physiological and molecular traits in AB211.

### Root exudates promote rhizosphere persistence

Genome-wide analysis focused on genes that exceeded a predefined log2 fold-change threshold (|log2FC| ≥ 1). However, several biologically important colonization-associated genes exhibited statistically significant, but moderate fold changes. Hence, representative regulatory genes that undergo subtle transcriptional modulation were selected for real-time expression analysis. For example, the density-dependent maintenance of bacterial populations, also known as quorum sensing, has been enhanced by treatment with root exudates. The S-ribosylhomocysteine lyase (*lux*S, BCV52_04890), involved in the quorum-sensing pathway, regulates bacterial multicellular communication and behavior^37^. It showed a 0.8-fold (relative fold change) upregulation in the transcriptome analysis, yet exhibited a 3-fold upregulation in real-time expression, suggesting that root exudates induce the quorum-sensing ability of AB211 to maintain its population during the early phase of the interaction (Supplementary Table S4, Supplementary Fig. 5A).

Initial attachment to the root surface is essential for the establishment and persistence of PGPR in the rhizosphere. This process is often mediated through biofilm formation by PGPR. Biofilms are multicellular structures enveloped by a polysaccharide matrix and physically attached to plant surfaces via surface-attachment appendages such as pili and flagella^38^. Biofilm formation is regulated by complex molecular mechanisms, including extracellular polymeric substance (EPS) production, adherence to the substrate surface, and expression of sporulation-related transcriptional regulators. The Type IV pilus system critically influences twitching motility of PGPR, thereby establishing strong root colonization through adhesion^39^. Profound upregulation of type IV pilus ATPase (*com*GA, BCV52_06740) and assembly protein (*com*GB, BCV52_06745) during the stationary phase indicated that the root exudates exposure significantly induced the surface attachment potential of AB211 for successful root colonization during the stationary phase (Fig. 3D). strengthening root colonization incidents. Sporulation-related genes, such as *spo*0A (BCV52_06965) and stage V sporulation gene *spo*VG (BCV52_00990), are positive regulators of biofilm formation in the *Bacillus* group^40, 41, 42^. The significant upregulation of these genes in AB211 was observed by real-time PCR, with more than 2–fold increases in relative expression in response to root exudates (Supplementary Fig. 5A). Hence, root exudates remodeled the physiology of AB211 to enable successful colonization in the rhizosphere (Fig. 3E).

Plants produce toxic methylglyoxal as a by-product of various metabolic pathways under stressed conditions^43, 44^. To survive in the competitive rhizosphere, PGPR must employ specific mechanisms to overcome these stresses. The glyoxalase pathway is reported as a mechanism for microbial detoxification of methylglyoxal, converting it into D-lactate^45^. Several reports suggest that glyoxalase-producing PGPR isolates can effectively alleviate methylglyoxal stress in plants, thereby strengthening mechanisms of abiotic stress tolerance^46, 47^. The two-fold relative upregulation of glyoxalase genes in the exponential phase indicates that AB211 can degrade stress components in root exudates, reduce plant stress, and promote phytobeneficial functions (Supplementary Fig. 5B). As mentioned in Supplementary Fig. 1A, root exudates also contain diverse secondary metabolites, such as phenolic compounds and organic acids, that function as antimicrobials against rhizosphere-dwelling microbes^48^. Successful rhizosphere colonizers are favored by their ability to metabolize these organic chemicals through the expression of the phenolic acid decarboxylase (*pad*C, BCV52_02340) gene. Exposure to root exudates enhanced *pad*C expression in AB211 (Supplementary Fig. 5C), thereby increasing its potential to compete and persist in the rhizosphere. Similarly, upregulation of D-cysteine desulfhydrase (*dcy*D, BCV52_09590), involved in sulfur metabolism and H_2_S production, indicates enhanced protection in the oxidative-stressed rhizosphere environment^49^(Supplementary Fig. 5C).

### Proteomics changes mirrored transcriptomic reprogramming

To validate whether the transcriptome-level changes translated into corresponding alterations in protein abundance, we performed a global proteome analysis under similar conditions of root exudates supplementation. Quantitative proteomics revealed extensive alterations in protein abundance in AB211 following root exudates treatment (Fig. 4A). Differential abundance analysis revealed significant upregulation of 183 proteins during the exponential phase compared to the control group (log2 fold change > 0.5, p-value < 0.05) (Supplementary Table S5). In contrast, fewer proteins were significantly down-regulated (Fig. 4A). Several proteins associated with translation, metabolism, nutrient acquisition, and stress adaptation exhibited larger fold changes and strong statistical significance, indicating that root exudates induce coordinated remodeling of the bacterial proteome during the early stages of rhizosphere adaptation.

Gene ontology enrichment analysis of differentially expressed proteins revealed that the proteomic response was dominated by proteins associated with translation, ribosome functions, and nucleotide biosynthesis (Supplementary Fig. 6). The significantly up-regulated proteins were functionally associated with ribonucleotide biosynthesis (35.63%), carbon-nitrogen bond-forming ligase activity (18.39%), ribosome biogenesis and activities, isoprenoid metabolism, and GTPase activity (Fig. 4B). Root exudates exposure tightly regulated the small and large ribosomal subunits, rRNA, RNA-binding proteins, and ribonucleoprotein complexes. Elevated purine and pyrimidine biosynthesis and metabolic processes also support the stable synthesis of essential proteins (Fig. 4C). A higher abundance of proteins involved in amino acid and secondary metabolite biosynthesis, such as isoprenoids, reflects active metabolic remodeling (Fig. 4B & C). Hence, differential protein abundance analysis indicated widespread proteomic remodeling, with significant modifications in the translational machinery (Fig. 4C). Up-regulated proteins were predominantly associated with nucleotide biosynthesis and metabolism, protein synthesis, isoprenoid metabolism, and cellular growth. Hence, the proteomics response revealed extensive remodeling of the protein synthesis machinery and central metabolic pathways in AB211 in response to root exudates exposure (Fig. 4C).

### Evidence of multilayered molecular reprogramming of AB211 in response to root exudates

The integrated profile of up-regulated genes and their corresponding proteins uncovered that the key functions such as nutrient transport and acquisition, signal responses, secondary metabolite biosynthesis, protein synthesis, genome repair and maintenance, chemotaxis, amino acid biosynthesis and redox homeostasis exhibited consistent upregulation with strong statistical significance in response to root exudates (Fig.4D). Moreover, correlation values indicated that functional attributes of significantly up-regulated proteins were extensively remodelled by few transcriptional regulators like genes encoding DinB family proteins (BCV52_20700), L-cysteine binding proteins (tcyJ, BCV52_18570), GntR family proteins (BCV52_08545), SpxA proteins (BCV52_25235) (Supplementary Fig. 7). Overall, the integrated data suggest that root exudates significantly alter bacterial behavior during the exponential phase, strengthening beneficial crosstalk during the early phase of the interaction.

### Ecological performances of AB211 in nature

Both physiological and molecular investigations demonstrated that root exudates influence the coordinated reprogramming of AB211, thereby activating metabolic pathways associated with rhizosphere adaptation. Therefore, we hypothesized that this molecular reprogramming might enhance the ecological competence of AB211, enabling it to establish itself in the native maize rhizosphere and restructure the resident microbiome. To test this hypothesis, we conducted a pot experiment using field-conditioned soil harboring a naturally assembled, taxonomically diverse maize-associated microbial community from the previous growing season, thereby establishing an ecologically relevant system for evaluating PGPR recruitment and rhizosphere microbiome dynamics. AB211 was introduced into this field-conditioned maize soil at a low inoculum density (1 × 10^5^ CFU/ml), approximating indigenous microbial abundance, thereby providing a realistic ecological context for evaluating AB211’s capacity to invade and modulate the native maize rhizosphere microbiome. Three amendment treatments were evaluated:(1) buffer (0.1 X PBS) control, (2) AB211 inoculation, and (3) maize RE-primed AB211 inoculation. Amendment with AB211 or RE-primed AB211 promoted plant growth compared with the untreated control, showing significant differences in root and shoot length throughout 21-days (Supplementary Fig. 8A). Although AB211 promote plant growth even in the presence of native soil microbiome, inoculation with RE-primed AB211 demonstrated a distinct phenotypic response, where a modest but consistent enhancement in plant root growth compared with the buffer control from 7 days onward were observed (Supplementary Fig. 8B-C). The observed developmental pattern indicates that the ecological competence of AB211 is intimately linked to rhizosphere developmental stages.

### AB211 restructures the native maize rhizosphere microbiome

To assess the impact of AB211 on the assembly of the maize rhizosphere microbiome, we employed complementary 16S rRNA gene amplicon sequencing and shotgun whole-metagenome sequencing. Soils collected immediately after amendment (0day), before seed sowing, were designated as bulk soil. In contrast, soil samples collected after seed sowing were classified as rhizosphere soil due to the continuous influence of root exudation following germination and plant development. The 16S rDNA amplicon sequencing revealed a shift in microbiome diversity upon AB211 amendment of the native rhizosphere soil microbiome (Fig. 5). The comparable Shannon diversity indices observed at day 0 indicate that the initial inoculation of AB211represented only a minor fraction of the native soil microbial community. Hence, the alpha diversity of the bulk soil microbiome remained unaffected (Fig. 5A). In the rhizosphere, the amended soils exhibited a gradual increase in alpha diversity beginning at 14 days, suggesting a gradual restructuring of the microbial community. However, only a marginal effect was observed at 21 days (Fig. 5A). Compositional analysis revealed that the maize rhizosphere was dominated by Actinobacteriota, Acidobacteriota, Chloroflexi, Bacteroidota, Firmicutes, and Proteobacteriota (Fig. 5B). This observation was reflected in the diversity abundance profile, where rhizosphere soils exhibited reduced microbial diversity relative to bulk soil, along with selective enrichment of several taxa like *Bacillus, Fictibacillus,* agricultural soil bacterium SC-I-84 and *Luteolibacter* in rhizosphere soil, hereby demonstrating the rhizosphere effect phenomenon (Fig. 5C). Estimation of the relative abundance across samples suggested that actinobacteriota, chloroflexi, bacteroidota, and firmicutes have enriched following amendment, particularly at14 days and 21 days (Supplementary Fig. 9). However, at 7 days, the abundance of actinobacteriota and acidobacteriota was significantly reduced in AB211-amended soil with unaltered abundance of proteobatceriota (Supplementary Fig. 9A, B &F). On the contrary, the RE-primed AB211 amendment resulted in a significant increase in the abundance of actinobacteriota and proteobatceriota at the same time point suggesting that root exudates priming modulates AB211’s ability to influence resident microbial community (Supplementary Fig. 9A&F).

**Fig. 5.**
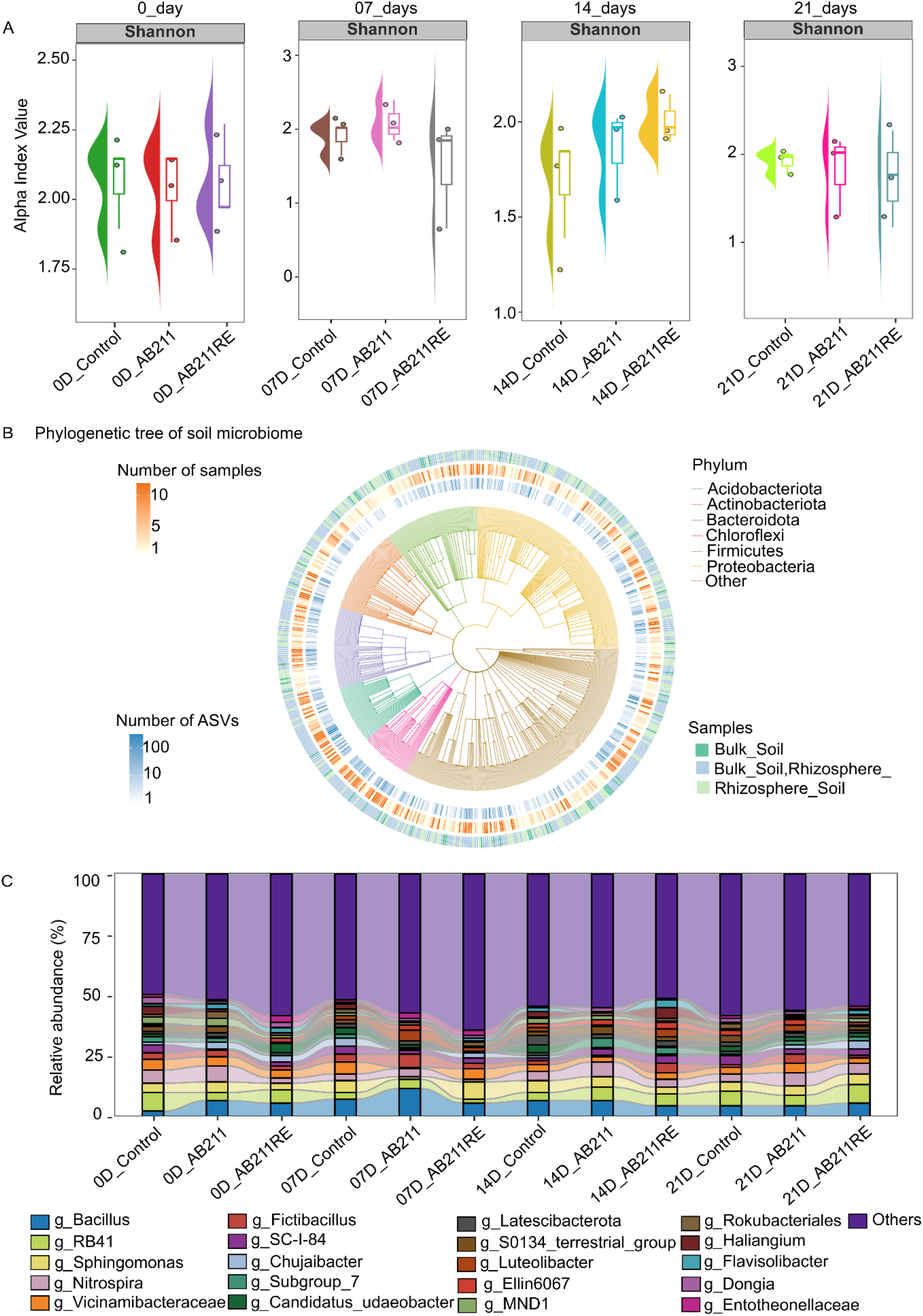
Alpha diversity of maize rhizosphere microbiome at 0 day, 7 days, 14 days, and 21 days upon amendment with buffer (control), AB211, and root exudates (RE) primed AB211. (A). Phylogeneetic tree generated from metataxomic diversity (B). Metataxonomic-based relative abundance of the top 20 dominant taxa in amended soil (C).

### AB211 selectively recruits beneficial microbial taxa

In-depth understanding of compositional variation within each microbial group using shotgun metagenomics revealed a nearly identical diversity pattern across the amended rhizosphere soils. Among the top twelve identified species, beneficial PGPR such as *Priestia aryabhattai*, *Priestia megaterium*, *Streptomyces nigra*, and *Bradyrhizobium* sp. CCBAU 51765*, Methylobacterium nodulans* exhibited profoundly stable enrichment in AB211 amended soil (Fig. 6A). The abundance profile of Actinomycetota group revealed that the relative abundance of *Streptomyces griseorubiginosus and Prescotella equi* have enriched in amended soil during early stages of interaction (07_days) (Fig. 6B).

**Fig. 6.**
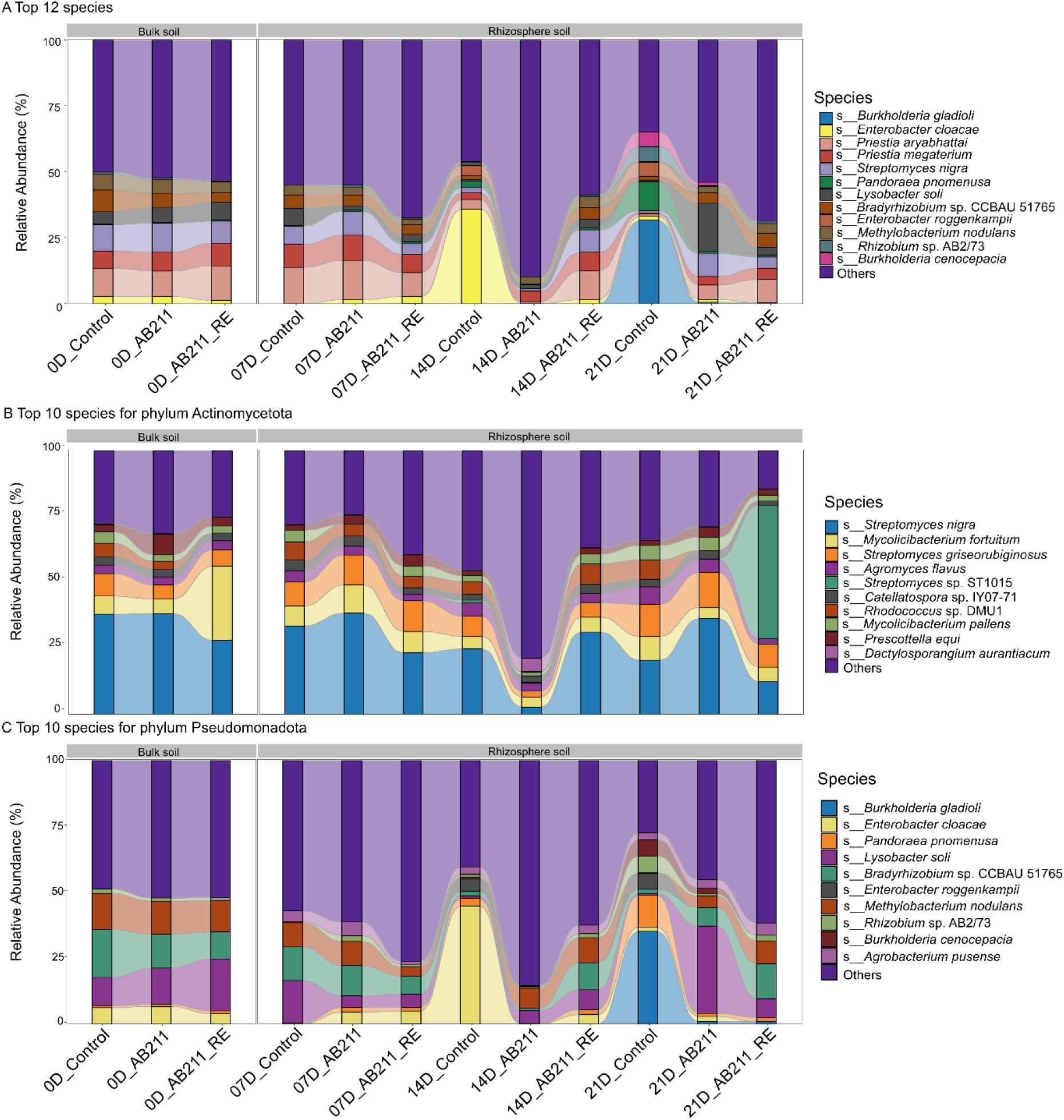
Taxonomic diversity as revealed from shotgun metagenomics. (A) representing the top 12 enriched species, following enrichment of the top 10 species of Actinomycetota (B) and Pseudomonadota (C).

Notably, AB211 amendment has enriched antimicrobial producing-*Dactylosporangium aurantiacum* at day 14, resulting in reduced abundance profile compared to control (Fig. 6B). *Agromyces flavus*, *Catellatospora* sp. IY07-71 decreased in amended soil with a surprising enrichment of *Streptomyces* sp. ST1015 in RE-primed AB211 amended soil at day 21 (Fig. 6B). Amendment with AB211 and RE-primed AB211 resulted in early recruitment of *Enterobacter cloaceae, Pandoraea pnomenusa,* and *Rhizobium* sp. AB2/73 with respect to control soil at day 7, indicating the ability of AB211, whether or not primed with root exudates, to selectively recruit beneficial bacteria during early stages of plant development (Fig. 6A & C). Compared with the control, the dominant prevalence was *Lysobacter soli* and *Bradyrhizobium* sp. CCBAU 51765 and *Methylobacterium nodulans* in the amended soil, especially under RE-primed conditions, indicate AB211’s ability to reshape a healthy rhizosphere by selectively enriching beneficial species within a taxonomically diverse microbiome (Fig. 6C).

### AB211 is established within the core maize rhizosphere microbiome

Metataxonomic analysis revealed the dominant appearance of Firmicutes in amended soil. Selective screening of the 16S rDNA amplicon for *Priestia aryabhattai* AB211 using nucleotide BLAST against closely matched representative sequences revealed a 100% match for two OTUs to the reference, identifying them as representatives of AB211. The relative abundance of these OTUs remained consistently high in amended rhizosphere soil, with a gradual increase from 14 days onwards (Supplementary Fig. 10A). This diversity pattern was also evident in the shotgun metagenomic distribution of Bacillota, showing the persistence of *Priestia aryabhattai* and *P. megaterium* in rhizosphere soil. Although diversity was severely altered, with reduced abundance of *P. aryabhattai* at 14 days, the rhizosphere was enriched with *P. megater*ium and *Neobacillus* sp. SuZ13*, Rossellomorea marisflavi,* and *Bacillus pseudomycoides*, indicating the efficacy of AB211 to recruit plant beneficial microbes in the rhizosphere (Supplementary Fig. 10B). The overall diversity profile uncovered the potential of AB211 to efficiently compete with the native core microbiome of the maize rhizosphere and establish itself as one of them, while fostering colonization by beneficial microbes functioning as keystone species. Hence, the modulation in plant development was not only the result of AB211-mediated enrichment of beneficial microbes but also due to the successful colonization of AB211 itself, which exhibits phyto-beneficial traits.

### AB211 rewires microbial communication network

Enrichment of microbial functions was predicted from the amplicon sequences using PICRUSt2. Functional profiling of the rhizosphere microbiome revealed that AB211 amendment promoted the enrichment of microbial functions associated with plant growth promotion, nutrient cycling, and ecological adaptation in maize. Most biological pathways, such as nitrogen metabolism, carbon fixation, tryptophan-associated functions, chemotaxis, and siderophore-associated functions, were enhanced in AB211 and RE-primed AB211-amended rhizosphere soil (Fig. 7A). The abundance of pathways related to chemotaxis-associated functions at 7days suggests that the AB211 amendment has influenced the early root exudation pattern and composition, thereby facilitating the recruitment of microbes in the rhizosphere. Additionally, induction of carbon– and tryptophan-associated pathways could correlate with the utilization of root exudates nutrients by recruited microbes, indicating ongoing multitrophic interactions in the rhizosphere. Interestingly, siderophore-associated functions have been comparatively more stimulated in AB211-amended soil than in the RE-primed AB211-amended soil, both at day 7 and day 14. The predicted enrichment of community-level functions suggests that the AB211 amendment may facilitate the establishment of a functionally resilient rhizosphere microbiome, thereby improving soil health and supporting plant growth and development.

**Fig. 7.**
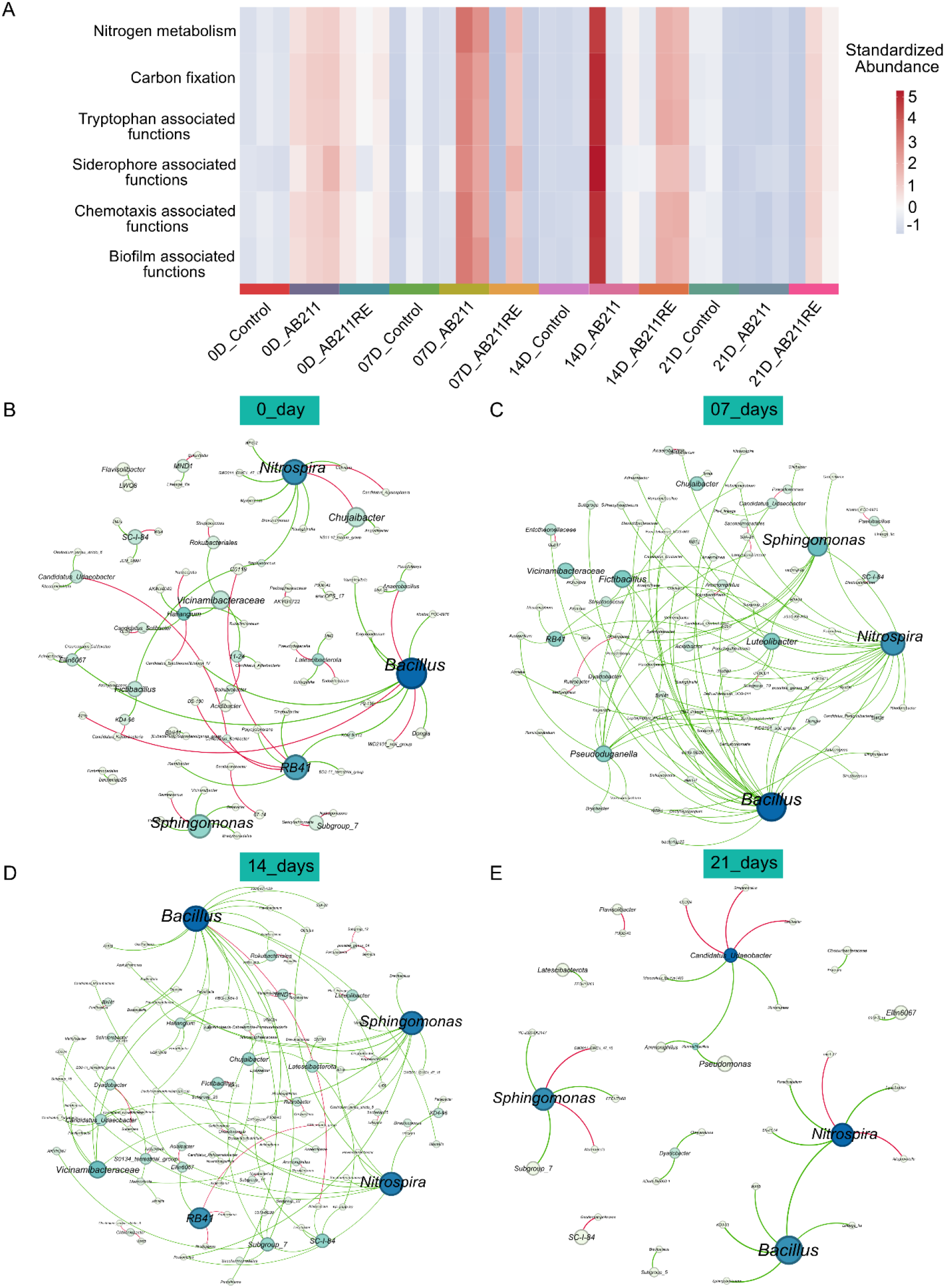
Community-level functional enrichment and interactions in amended soil using PiCRUSt2. The heatmap shows the predicted KEGG pathway abundances across samples. The color intensity indicates the normalized Z-score of the KEGG pathway abundances (A). The microbiome interactome network reveals shifts in microbial communication and interactions at 0 day (B), 7 days (C), 14 days (D), and 21 days (E). The network was generated using the SparCC algorithm with a 0.6 correlation cutoff and a p-value < 0.05 (FDR adjusted).

To investigate the driving factors behind such community functions, microbial network analysis was performed with the top microbial taxa. The microbial network of the maize rhizosphere was dominated by *Bacillus*, *Sphingomonas*, and *Nitrospira* across four sampling days, including bulk and rhizosphere samples, indicating that these three genera were positively correlated with most other beneficial genera, thereby stabilizing a healthier soil microbiome (Fig. 7B-E). Positive and complex interactions of these organisms with several phyto-beneficial microbes, such as *Rhizobia*, *Azospirillum*, *Brevundimonas*, *Burkholderia*, and *Luteolibacter* over 7 and 14 days, indicate root exudates-mediated selection of rhizosphere microbes, thereby strengthening a core healthy rhizo-microbiome for better plant growth and development (Fig. 7C and D). These observations indicate that the reintroduction of AB211 has been associated with the enrichment of beneficial microbial populations, thereby modulating the maize rhizo-microbiome. Conversely, by day 21, the microbial interaction network exhibited reduced complexity, indicating a transition toward a more stabilized community structure. The dominant taxa influenced a smaller subset of interacting microbial groups, hence maintaining a steady ecological niche (Fig. 7E). Thus, by the 21-day seedling stage, the maize plant appears to have established a relatively stable core rhizosphere microbiome.

### AB211 remodels the functional landscape of the maize rhizosphere microbiome

Predictive information on community-level functional turnover alone is insufficient to guide field-level applications aimed at developing sustainable agricultural strategies to enhance plant resilience. Developing an effective biofertilizer formulation requires a deeper understanding of the key microbial species and their interactions that drive beneficial plant–microbiome associations. To understand the capacity of enriched beneficial species involved in robust reconstruction of metabolic pathways of soil, we correlated the bacterial enrichment patterns with their beneficial functional traits (Fig. 8). Notably, metagenomics provided clear insight into the underground crosstalk mechanism, revealing the benefits of RE priming. Gene ontology-based pathway enrichment analysis coupled with species identification revealed enrichment patterns of nutrient utilization and metabolic pathways, such as nitrogen and carbon fixation. We observed that *Bacillus aryabhattai* (also referred to as *Priestia aryabhattai* due to database mismatches) predominantly regulated nutrient cycling in the rhizosphere. Mainly, these fixation pathways were selectively enriched in the RE-primed AB211-amended rhizosphere at 7 and 14days (Fig. 8A-B). The increase in the nitrogen-fixation pathway in the 14-day RE-primed AB211-amended soil could be correlated with the association of *Rhizobia*, including *Bradyrhizobium japonicum, B. liaoningense, Azospirillum oryzae, and Azoarcus* sp. KH32C as observed in the species-level abundance profile (Fig. 8A).

**Fig. 8.**
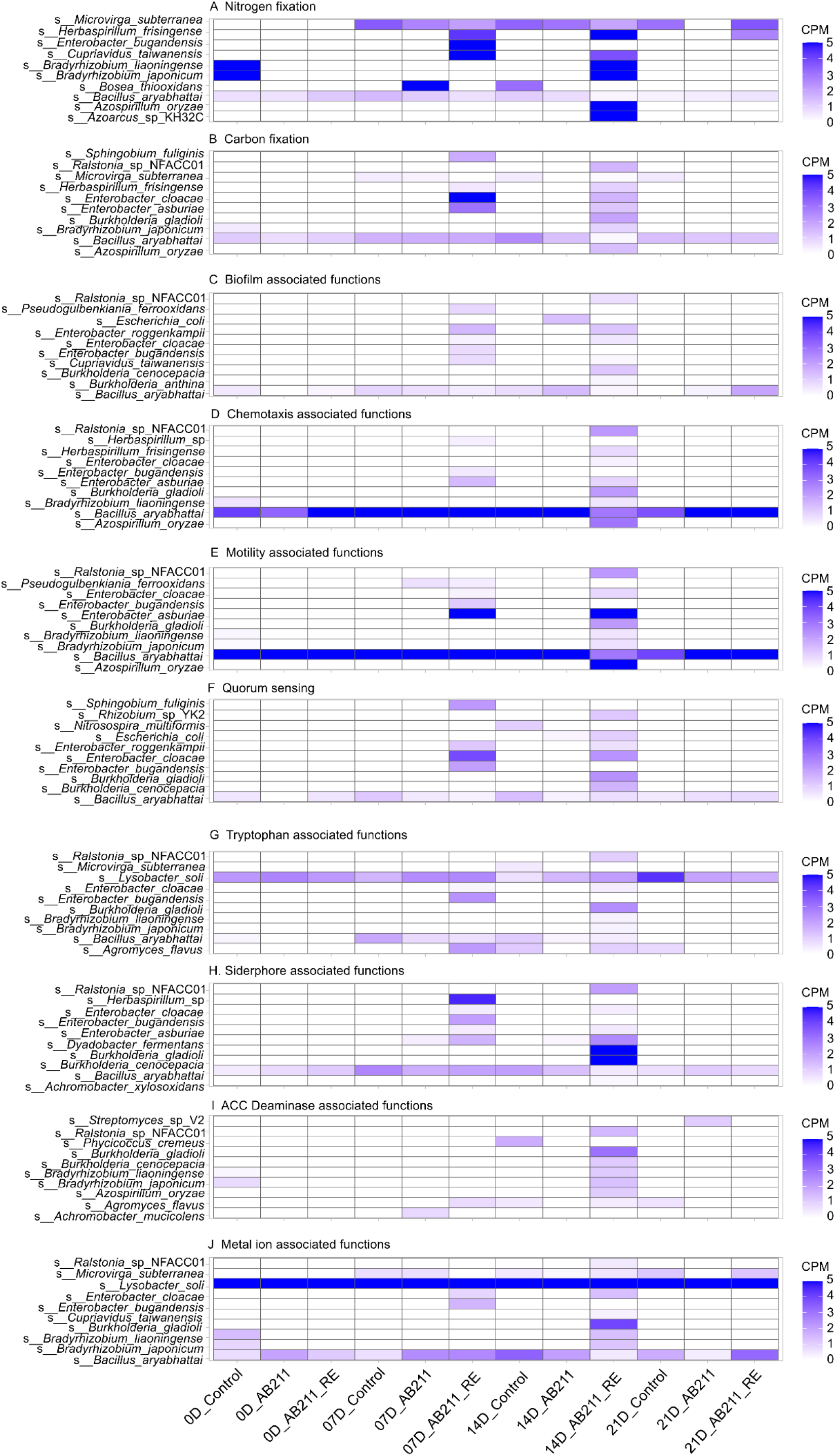
Species-level resolution of enrichment of beneficial functional traits in amended soil. The heatmap shows the microbial pathways obtained from HUMANn across samples. Color intensity represents the normalized counts-per-million (CPM) values.

Similar functional enrichment was observed in plant-microbe interaction-related pathways—biofilm formation, chemotaxis, motility, and quorum sensing (Fig. 8C-F). *Bacillus aryabhattai* (*Priestia aryabhattai*) was consistently detected in all pathways. The quorum-sensing and chemotaxis-related pathways have been significantly increased on the 7th and 14th days of RE-primed AB211-amended rhizosphere soil.

Further investigation of plant beneficial properties of the identified top bacterial species revealed enhanced modulation of tryptophan-associated function in the rhizosphere soil on the 7th and 14th days, with the highest increase in RE-primed AB211-amended soil (Fig. 8G). Both siderophore and ACC deaminase activity have been enhanced in the amended rhizosphere, following a similar pattern (Fig. 8H-I). Functional enrichment of metal-ion degradation, adsorption, bioconversion, etc., in RE-primed AB211-amended soil was observed. Therefore, root exudates priming reprogrammed AB211 at molecular level, The overall functional enrichment pattern suggests that RE priming has efficiently reprogrammed AB211 at molecular level, potentially enhancing its capacity to reshape rhizosphere microbiome, enriching phytobeneficial functions in rhizosphere, and ultimately contributing to improved maize growth.

## Discussions

Declining soil quality and extensive usage of agricultural fields have intensified the need for sustainable strategies to harness beneficial plant-microbe interactions. In this context, the rhizosphere soil serves as an active nexus of multitrophic biological interactions, dynamically influenced by metabolites released from living organisms ^50^. Root exudates, in particular, are recognized as one of the major determinants of microbial recruitment in the rhizosphere. The nutrient-rich property of root exudates facilitates microbial adaptation in the rhizosphere ^6^. The exudation pattern is also highly dynamic across plant developmental stages ^51^. Non-targeted metabolomic studies of maize root exudates have demonstrated that the early developmental stage is associated with increased sugar exudation. In contrast, at later stages, the release of phenolics and secondary metabolites is significantly enhanced ^52^. Our findings support such observations, confirming the presence of glucose, sucrose, amino acids and organic acids in root exudates (Fig. 2). Similar exudation patterns with increasing abundance of antimicrobial-like secondary metabolites like coumarin, (S)-N-methylcoumarin, copal-8-ol diphosphate etc., during 21 days of plant growth indicates plant’s ability to shape microbial assemblages depending on its growth stages (Supplementary Fig. 1A). However, how these metabolites influence the molecular mechanisms that enable beneficial bacteria to establish within, and subsequently influence native microbial community, remains poorly understood.

A previous study on the influence of root exudates on *Bacillus amyloliquefaciens* FZB42 showed extensive transcriptional modulation of bacterial metabolic pathways involved in plant-microbe interactions, such as nutrient metabolism, chemotaxis and motility, and the synthesis of antimicrobial peptides^16^. Additionally, the secretome profile of *B. amyloliquefaciens* FZB42 in response to maize root exudates revealed that proteins involved in nutrient utilization pathways, hydrolysis of oligosaccharides, chemotaxis and motility, and antioxidative activities, were secreted at higher levels, supporting the impact of root exudates on plant-microbe communication^53^. Furthermore, many studies have highlighted the differential ability of microbes to adapt to specific rhizospheric environments. For instance, a comparative study of eight *Pseudomonas* strains examined distinct responses to root exudates, showing marked differences in the expression patterns of genes involved in carbohydrate and amino acid metabolism, and copper and iron homeostasis across the strains tested^13^. These microbial responses indicate that prior exposure to root exudates influences bacterial adaptation to fluctuating rhizosphere deposits. However, previous studies have primarily focused on microbial responses to root exudates or the community-level effects of microbial inoculation on plant–microbe interactions. The extent to which root exudates-mediated bacterial preconditioning enhances the plant growth-promoting efficacy of PGPR, influences its rhizosphere competence, and reshapes the functional potential of native microbial communities in a heterologous soil environment remains largely unexplored. Hence, it is necessary to explore the rhizosphere adaptation strategies of beneficial microbes *in vitro* and in native soil, thereby bridging the gap between laboratory research and field applications.

Here, we addressed this research gap by investigating how maize root exudates influence the physiology of *Priestia aryabhattai* AB211 and how root exudates-mediated preconditioning affects its interactions with the native microbial community in the maize rhizosphere. AB211, a potent plant growth-promoting bacterium originally isolated from the tea rhizosphere in Darjeeling, has been shown to promote maize growth under sterile conditions ^28^. We therefore explored its potential to improve plant– microbe interactions and inform sustainable agricultural practices. Transcriptomic and proteomic analyses revealed extensive physiological reprogramming in AB211 following exposure to root exudates, including changes in pathways associated with chemotaxis, motility, and siderophore production. These responses suggest that maize root exudates may enhance bacterial adaptation to the rhizosphere and strengthen traits relevant to plant growth promotion. Accordingly, we hypothesized that root exudates could influence AB211’s transition from a free-living soil microbial state to a rhizosphere-adapted colonizing state, even before it encountered the rhizosphere.

To investigate this hypothesis, we primed AB211 with maize root exudates and amended the maize rhizosphere soil, which is a non-native environment for the organism. Studies have shown that the native rhizosphere microbiome holds tremendous potential for supporting plant growth^21^. Whereas non-native PGPR such as *Bacillus subtilis* and *Pseudomonas fluorescens* function as potent microbes under sterile conditions, they face difficulties with rhizosphere colonization due to a lack of home-field advantages^21, 54^. Nevertheless, the functional legacy of the non-native microbes persists in the rhizosphere^55^. In the current study, heterologous introduction of AB211 into the maize rhizosphere selectively altered the existing diversity and abundance profiles of the native microbiome. The transient dominance of frequently beneficial taxa over 21 days, including *Bacillus*, *Sphingomonas*, *Nitrospira*, and Actinobacteria, suggests that microbiome assembly followed selective restructuring rather than generalized assembly. Hence, root exudates priming reflected a distinct influence on AB211, thereby governing the ecological selection pressure in the rhizosphere. A similar pattern of microbial enrichment has also been observed in numerous field applications of beneficial bacterial inoculation in natural field soils, suggesting a positive influence of PGPR amendment on the rhizosphere microbiome^18, 56^. Moreover, increased complexity of the rhizosphere microbial network during early seedling stages following AB211 amendment was associated with greater network connectivity and co-occurrence among microbial taxa. This pattern is consistent with enhanced ecological interactions of functionally complementary microbes, indicating the selective enrichment of essential microbes to support early plant growth. In contrast, a simplified network in later stages indicates that the rhizosphere microbiome stabilizes to meet the specific needs of mature seedlings. These observations are supported by recent models proposing that plants assemble a functionally optimized core microbiome through successive ecological filtering rather than indiscriminate microbial recruitment^9^.

Functional reconstruction revealed coordinated enrichment of pathways supporting soil health and plant growth in RE-primed AB211 amended soil, including nitrogen and carbon fixation, chemotaxis, quorum sensing, biofilm formation, biosynthesis of siderophores and tryptophan, ACC deaminase activity, and metal-ion-related processes. These functions broadly regulate nutrient acquisition, microbial communication, and stress adaptation, indicating that prior exposure to root exudates not only improved AB211’s ability to modulate microbiome diversity but also rewired the rhizosphere microbiome’s metabolic capacity. This observation correlated with microbial crosstalk within the maize rhizosphere microbiome in the presence of *Sphingomonas* sp. Hbc-6 inoculation, following a significant increase of metabolites mediating recruitment of beneficial microbes like *Variovorax, Comamonas, Methylobacterium,* and *Streptomyces*^57^. Subsequently, shotgun metagenomics further supported these observations of functional trends while providing species-level resolution, thereby strengthening the biological interpretation inferred by amplicon sequencing. Together, these observations support a model in which root exudates mediated the physiological reprogramming of AB211, which contributed to subsequent rhizosphere microbiome assembly.

Taken together, our findings suggest that root exudates serve as chemical signals that modulate the physiology and behavior of AB211. Prior exposure to root exudates simultaneously reprogrammed the molecular mechanisms of AB211, thereby potentially enhancing traits associated with rhizosphere competence under the native maize rhizosphere. This work extends the current understanding of beneficial plant-microbe communication by demonstrating that root exudates play a role beyond microbial recruitment in the rhizosphere, indirectly influencing microbiome assembly through the physiological reprogramming of non-native beneficial PGPR^58^. Collectively, these mechanistic insights provide a framework for rhizosphere microbiome engineering using ecologically adapted bioinoculants that can stably integrate into and reshape native soil microbial communities. This strategy holds considerable promise for developing resilient, microbiome-informed biofertilizer applications to enhance crop productivity and sustainability.

## 2. Materials and methods

### 2.1. Description of microbial inoculant and plant variety

In this study, we used AB211 as a model PGPR in all the experiments. AB211 was previously isolated from the tea rhizosphere of Darjeeling^28^. Draft genome sequencing revealed that AB211 is metabolically versatile and exhibits robust plant growth-promoting traits (https://www.ncbi.nlm.nih.gov/nuccore/MCAN00000000.1). AB211 was routinely cultured in minimal medium (M9) and incubated at 37 °C in a rotary shaker (150 rpm).

The Early Golden Bantam (EGB) variety of maize (*Zea mays*) was selected for all *in planta* experiments. Surface-sterilized maize seeds were imbibed in sterile water for 30 minutes and sown in sterile soil-rite. Three seeds were sown in each pot and incubated in a plant growth chamber at 25 °C under an optimized photoperiod of 16 h light and 8 h dark until the desired seedling stages were reached. Plants were regularly watered with tap water.

### 2.2. Extraction of root exudates

As described above, maize seedlings were cultivated for 7, 14, and 21 days. Maize seedlings were carefully loosened from the pots and soaked in a bucket of tap water for a few minutes, allowing partial removal of loose soilrite from the roots without causing physical stress. Then, the roots were gently agitated in a fresh tray of water, repeated for three to four times until the water remained clear of any soil debris ^59^. Twenty to twenty-two washed roots were immersed in 200 mL of sterile water in a beaker and incubated for 16 hours at 28 °C after being shaken at 146 rpm. Later, the water containing the root exudates was centrifuged (3,000 × g for 15 minutes), and the clear supernatant was freeze-dried to obtain the root exudates powder. The concentrated root exudates powder was weighed, resuspended in sterile water, and filtered through a 0.22 µm sterile syringe filter. For long-term storage, aliquots (100 µl) were kept at –80 °C.

### 2.3. Estimation of amino acid, organic acid, and carbon components in maize root exudates

The amino acid composition in root exudates was analyzed using HPLC (Shimadzu). The root exudates sample was pre-derivatized with o-phthaldialdehyde (OPA) and injected into the HPLC column (GROM-SIL OPA-3). Separation was achieved using a gradient of two solvents: solvent A [tetrahydrofuran (0.70 %) added phosphate buffer (25 mM, pH 7.4)] and solvent B [(a mixture of acetonitrile, methanol, and 25 mM phosphate buffer, pH 7.4 in a 15:35:50 ratio (v/v/v)], following Fan et al. 2012, with slight modifications^16^. Fluorescence intensity of the derivatized amino acids was detected with excitation at 330 nm and emission at 450 nm.

Ion chromatography was performed to quantify organic acids in RE using a Dionex IonPac AS 11-HC column, following a KOH concentration gradient from 10 mM to 80 mM^16^. Known standards were used to identify the organic acids by comparing their retention times.

Sugars were analyzed using a GC-MS (Thermo Fisher, USA). An aliquot of 100 μl RE was derivatized by reconstituting in 50 μl of methoxyamine hydrochloride in anhydrous pyridine and kept at 37 °C for 2.5 hours, followed by a 40-minute reaction with 50 μl of N-methyl-N-trifluoroacetamide at the same temperature. Subsequently, 1 μl of the derivatized sample was injected into the GC column ^16^.

#### Untargeted profiling of maize root exudates

The root exudates profile was assessed using an Xevo® G2-XS Q-Tof MS equipped with a Waters ACQUITY UPLC M-Class System (Waters Corporation, Milford, MA, USA). 20 µl root exudates (10 mg/mL) were diluted with 40 µl formic acid (0.1%), and 20 µl of the diluted sample was aliquoted into vials for LC-MS/MS analysis. Details of the chromatographic separation are provided in Supplementary file 1. The raw data were processed using Progenesis QI software, following ChemSpider, KEGG, and PlantCyc as reference databases. Commonly abundant metabolites were selected by manual screening of the 7, 14, and 21-day root exudates profiles. The change in exudation pattern was visualized by plotting z-scores for the selected compounds.

#### Planktonic lifestyle of AB211 upon root exudates exposure

Overnight culture of AB211 was diluted to 0.05 (at OD_600_) and inoculated in complete M9 minimal medium broth (Amresco). Previously extracted, filter-sterilized maize root exudates were added to the culture at concentrations of 0.25 mg/ml and 0.5 mg/ml. The untreated culture was maintained as a control. The culture was incubated at 37 °C with shaking (146 rpm), and turbidity was measured at 30-minute intervals at 600 nm. Finally, the results were plotted to obtain the growth pattern. The methods for in vitro plant growth promotion assays upon exposure to root exudates have been described in the supplementary methods.

#### Preparation of AB211 samples for RNA-Seq

A fresh culture of AB211 was inoculated into RE-supplemented (0.25 mg/ml) M9 minimal medium. The culture was incubated with shaking (146 rpm) at 37 °C. The cells were harvested at the exponential phase (O.D._600_ 1.0) and the stationary phase (O.D._600_ 2.0). Culture grown in M9 medium without root exudates supplement was considered the control. After centrifugation, the pellet was washed with fresh M9 medium. The resulting pellet was stored at –80 °C in RNA Later solution (Thermo Fisher Scientific) until further processing. Paired-end RNA-Seq was performed on the NovaSeq6000 150PE Illumina platform.

#### RNA-Seq analysis

The quality of the raw reads was assessed using FastQC v. 0.12.1 (with default parameters). Removal of adapter sequences was done with fastp_v.0.20.1, and the quality of the trimmed reads was re-evaluated with FastQC_v.0.12.1. High-quality filtered reads were indexed and aligned with *P. aryabhattai* AB211 (GCF_001858395.1_ASM185839v1) as the reference genome using Bowtie2 (V_2.4.4). The resulting alignment file (.bam format) was used for gene-level quantification with htseq-count (v.2.0.9). Differential gene expression was analyzed with DESeq2. The gene counts were used as input to DESeq2 for downstream statistical analyses to calculate log2 fold changes in treatment compared to control. Internal normalization and dispersion estimation methods were applied for statistical analysis. Throughout the manuscript, we have used p-value ≤ 0.05 and log2 fold change ±1 as cutoff values for significant differential gene expression. The downstream analysis was performed in R. The functional categories of differentially expressed genes were analyzed using eggNOGmapper and KEGG.

#### Quantitative real-time PCR

Total RNA was isolated from fresh cultures of AB211, supplemented with or without root exudates, using the two-step TRIzol method, following DNase I treatment. The first-strand cDNA was synthesized using the PrimeScript™ RT reagent Kit (Takara Bio, Japan) according to the manufacturer’s instructions, with random hexamers as primers. Primers for selected genes were designed using NetPrimer, Clone Manager, and Integrated DNA Technologies (IDT) software with standard parameters (Supplementary Table S6). SYBR Green Master Mix (TAKARA) was used for real-time PCR on a BioRad instrument. Primer specificity was confirmed by melt curve analysis and test PCR. Relative expression of the gene of interest was evaluated using the threshold cycle (Ct) values (2^-ΔΔCt) to determine the relative fold change^12^. Genes encoding *Lac*I, 50S ribosomal protein L18, and 16S rRNA were considered internal controls based on their non-significant expression levels in global transcriptomic datasets.

#### Proteome profile of AB211 upon root exudates treatment

Root exudates-treated AB211 cells were harvested during the exponential and stationary phases, and total protein was extracted using the ultrasonication method. The samples were run on SDS-PAGE for 10 minutes as part of downstream processing. 1×1 mm gel cubes were excised and serially washed, then reduced, alkylated, and digested with trypsin^60^. In brief, gel pieces were destained by washing with 300-800 µl of 70 % ammonium bicarbonate (ABC, 50 mM) + 30 % acetonitrile for 60 minutes, depending on the intensity of Coomassie stain and the gel size. The cycle of shrinkage and swelling was performed twice using acetonitrile and 50 mM ABC. Vacuum-dried gel pieces were sequentially incubated with 10 mM DTT and 55 mM iodoacetamide to reduce and alkylate protein samples. Then dried gel pieces were suspended in 15-200 µl of 15 ng/µl trypsin (Promega) dissolved in 25 mM ABC and incubated overnight at 37 °C. Protein was extracted from gel pieces by vortexing in 100-800 µl of formic acid (5 %) and acetonitrile (30 %) and sonicated for 5 minutes. Finally, the extracted protein sample was desalted using C18 desalting tips, and the digested peptides were loaded onto an Easy Spray Column PepMap^TM^ RSLC C18 (3 µm, 100 Å, 75 µm x 15 cm). Samples were analyzed on a Q-Exactive mass spectrometer coupled to a nano-flow HPLC system (Easy-nLC 1200, Thermo Scientific). Samples were eluted using a 60-minute gradient from 0.2 % formic acid in 5 % acetonitrile-water (solution A) to 90 % acetonitrile-water (solution B), with a flow rate of 300 nl/min.

#### Processing of proteomics data

The LC-MS (.raw) data (n=12) (including exponential and stationary phases with control groups) were converted to a Mascot-readable format (.mzml) using Proteowizard MSConvert (https://proteowizard.sourceforge.io/download.html) for downstream analysis ^61^. Mascot server 2.6 (https://www.matrixscience.com/) was used for MS/MS peak identification and characterization, with peptide tolerance of 1.2, MS/MS tolerance of 0.6, and a missed cleavage of 1 ^62^. The proteins from *Pristetia aryabhattai* (GCA_001858395.1_ASM185839v1) were used as a reference database. Protein identifications were accepted at less than 1% false discovery rate (FDR). The. dat files were uploaded to Scaffold 4 software (https://www.proteomesoftware.com/) to quantify proteins using specific parameters, including a protein threshold of 99%, a minimum of 2 unique peptides, and a peptide threshold of 95%, thereby generating a matrix of total spectral counts^63^. The raw protein matrix data were normalized (median and log2-transformed) and used to identify differentially abundant proteins between groups using limma, with log2 fold change > 0.5 and p-value < 0.05 ^64^.

#### Integration of proteomics and transcriptomics datasets

The differentially up-regulated proteins in the exponential phase compared to control groups (n=183) were mapped to their corresponding genes in the transcriptome dataset. Next, we used the supervised N-integration method DIABLO (Data Integration Analysis for Biomarker Discovery using Latent Variable Approaches for Omics Studies) from the MixOmics R package (version 6.32.0) to integrate proteomics and transcriptomics datasets^65, 66^. DIABLO with multiblock (s)PLS-DA was used to identify correlations (cut-off 0.7) between proteins and genes in the exponential phase.

#### Collection and preparation of field-conditioned native rhizosphere soil from field-grown maize

Rhizosphere soil was collected from field-grown maize (Zea mays cv. Early Golden Bantam) plants at a depth of 15–20 cm during April–May 2025 from the Madhyamgram campus of the Bose Institute, Kolkata. Plant debris, roots, and coarse particles were removed, after which the soil was air-loosened, gently crumbled by hand while wearing sterile gloves, and passed through a 2-mm sterile sieve to obtain a homogeneous and well-aerated soil matrix. Throughout the processing, care was taken to minimize external contamination. The processed soil was designated field-conditioned soil because it naturally harbored the indigenous core rhizosphere microbial community associated with field-grown Early Golden Bantam maize plants.

Fresh culture of AB211 was inoculated into minimal medium (M9), with or without a maize root exudates supplement from 14-day-old maize seedlings, at 0.25 mg/ml. The cells were harvested by centrifugation at 5,000 × g for 10 minutes, at an OD_600_ of 1.0 (3 × 10^8^ CFU/ml), i.e., the exponential phase. The harvested AB211 cells were resuspended in 0.1X PBS. 6 kg of field-conditioned soil was homogeneously mixed with the AB211 suspension (O.D._600_ of 0.1), maintaining an initial inoculum of 1 × 10^5^ CFU/ml. Now, 500 g of this conditioned soil was distributed among pots and used to sow maize (EGB) seeds and for further experiments.

Three seeds were sown in each pot conditioned with (1) AB211, (2) root exudates-primed (RE-primed) AB211, and (3) PBS or buffer control. Plant parameters, such as root and shoot length, were recorded at 7, 14, and 21 days. Rhizosphere soil samples from each treatment were also collected at a similar time interval. The plants were carefully uprooted, and loosely adhering bulk soil was gently removed by shaking. The soil tightly attached to the root surface, considered rhizosphere soil, was collected into tubes by scraping with a sterile spatula. Total soil DNA was extracted using the NucleoSpin Soil DNA Isolation Kit. DNA was also extracted from soil immediately after collection from the field, considered field soil, i.e., devoid of a fresh rhizosphere effect. The soil DNA samples were processed for amplicon and whole-genome metagenomic (WGS) analysis by Neuberg Diagnostics. Paired-end sequencing was performed (150 bp x 2) on the Illumina Novaseq6000 platform.

Amplicon sequencing was performed by targeting the V3-V4 region using V3_F (5’-CCTAYGGGRBGCASCAG-3’) and V4_R (5’-GGACTACNNGGGTATCTAAT-3’) primer pairs. In amplicon sequencing, library preparation was carried out using the Ultra II DNA Library Prep Kit for Illumina. The KAPA HyperPlus Kit was used to prepare libraries for shotgun metagenomic sequencing. A Qubit 4.0 fluorometer (Thermo Fisher Scientific) was used to quantify libraries using the DNA HS assay kit (Thermo Fisher Scientific). The library size was selected on the LabChip® GX Touch/GXII Touch using the DNA NGS 3K Assay.

#### Metataxonomic analysis

Microbiome diversity analysis was performed using QIIME2 (version 2024.10), with modifications^67,68^. Trimmed quality reads were denoised using the DADA2 algorithm^69^. Feature table and representative sequences were extracted using denoised and dereplicated sequences. Meta-taxonomic classification was performed using the trained SILVA 138.1 database to retrieve SSURef NR99 sequences^70^. The feature tables were exported in TSV format using the biom-convert tool. The artifacts (.qza) from QIIME2, including operational taxonomic unit (OTU) tables, taxonomy, phylogenetic trees, and metadata files, were imported and analyzed in R version 4.4.1 (2024-06-14 ucrt). The microbial diversity analysis was conducted using the MicrobiotaProcess R package^71^. Sequencing depth was standardized by performing rarefaction with the mp_rrarefy function, followed by the calculation of rarefaction curves (mp_cal_rarecurve function). The Shannon index of alpha diversity, and relative abundance of taxa were calculated (mp_cal_alpha and mp_cal_abundance function, respectively)^68^. Abundances of phyla across soil samples from different seedling stages were estimated from relative abundances, and the results were plotted (mp_plot_abundance function).

#### Phylogenetic tree of amplicon data

In 16S-Amplicon sequencing data, the taxonomic tree was generated by aggregating taxa to the genus level using treeio (version 1.32.0)^72^ and then plotted with the ggtree package (version 3.16.3)^73^. The raw data were normalized to relative abundance (%) to represent the top-abundant phyla and genera in each group, and the results were plotted using MicrobiotaProcess^71^.

#### Microbial network in amplicon data

The co-occurrence network of microbial genera was generated using the Sparcc algorithm in the microeco (version 1.16.0) R package^74, 75^. The samples from each day (0-day, 7-day, 14-day, and 21-day) were merged to obtain a day-wise overview of the interactomes. The correlation cutoff of 0.6 and p-value of < 0.05 were chosen to filter out low and non-significant interactions. The green and red edges in the microbial interactomes indicate positive and negative interactions, respectively, and the nodes represent microbes, with node sizes proportional to their relative abundances.

#### Predictive functional analysis

The Phylogenetic Investigation of Communities by Reconstruction of Unobserved States (PICRUSt2 – v2.5.2) was used to make metagenomic predictions based on 16S rRNA amplicon data to obtain the KEGG Orthologs (KO) ids and Metacyc pathways and their corresponding abundances^76^. The differentially abundant pathways were identified using the ALDEx2-Kruskal-Wallis test in the ggpicrust2 package (v.2.5.6) with p-values < 0.05^77^.

#### *In silico* tracking of AB211

From the QIIME2 taxonomy table, five OTUs representing the genus *Bacillus* were extracted along with their Feature IDs. The selected feature IDs were used to retrieve their corresponding DNA sequences from the representative table (representative.qza). The V3 and V4 regions of the 16S rDNA of AB211 were extracted from the full-length 16S sequence using the same V3_F and V4_R primer pairs as used in amplicon sequencing. NCBI nucleotide BLAST was performed among the five representative sequences using the AB211 V3_V4 sequence as a reference. The mean value was plotted, with 0-day soil samples represented as bulk soil and 7, 14, and 21-day samples as rhizosphere soil.

#### Analysis of shotgun metagenomics

The raw shotgun metagenomics data (.fastq) were processed for quality control using FastQC (Andrews 2023). Better-quality reads were filtered using Trimmomatic with the default parameters^78^. To remove host DNA sequences, filtered reads were processed with Kneaddata (Version 0.12.0) using the Maize genome as the reference. The host removed clean reads were used for taxonomy and pathway analysis using Kraken2 and Humann. R version (v_4.4.2) was used to plot the observations.

#### Taxonomic analysis and functional enrichment

Taxonomic assignment was performed using Kraken2 (version 2.1.1) with a confidence score of 0.1. Prebuilt RefSeq indexes: PlusPF (standard plus protozoa and fungi) (Version: 2024-1-12 – Downloaded: 2024–07-15T185612Z) were used to map reads to microbes^79^. Bracken (version 2.1) (with the reference database Prebuilt Refseq indexes: PlusPF (Version: 2024-01-12, k-mer: 35, Read: 100)) was used to re-estimate species-level abundance from Kraken2 reports^79^. Kraken-style reports from bracken were merged with the metadata file to produce a Biological Observation Matrix (BIOM) file using the kraken-biom tool^80^.

Here, we have used HUMAnN 3 (version 3.9) to profile microbial functional features (pathways and genes) on the European Galaxy server ^81, 82^. Briefly, whole metagenomic sequencing (WMS) reads were mapped to clade-specific markers to identify microbes using a reference database (mpa_Jun23_CHOCOPhlAnSGB_202403). Nucleotide-level mapping was conducted using the full ChocoPhlAn database (2023), and the translated search was performed using the full UniRef90 (2023) database in HUMAnN. The gene list was then mapped to Gene Ontology using the “Regroup” function. The data was normalized to counts per million (CPM) by the “Renormalize” function. Downstream analysis was performed using R.

## Statistical analysis

The significance of the growth curve was analyzed using two-way ANOVA, followed by Tukey’s multiple comparisons post hoc test, with the growth pattern of untreated AB211 as the control. Similarly, one-way ANOVA followed by Dunnett’s multiple comparisons test was performed to assess the significance of gene expression in AB211 upon treatment with root exudates, with untreated AB211 from the exponential and stationary phases as respective controls. For plant growth parameters, two-way ANOVA, followed by Tukey’s multiple comparisons test, was performed to assess significance in response to the amendment.

## Acknowledgment

This work was supported by an extramural grant [CRG/2018/001732] from the Science and Engineering Research Board (SERB), Government of India, and an intramural research grant from Bose Institute, Kolkata. Sangita Mondal was supported by a Senior Research Scholarship (Ref. 191620161234) from the University Grants Commission (UGC), India; Dhruba Bhattacharya was supported by the National Postdoctoral Fellowship (SERB-NPDF) [Ref. PDF/2021/003080] from SERB, Government of India; and Sriparna Mondal was supported by a Junior Research Fellowship from the University Grants Commission (UGC), India. The authors would like to acknowledge the Central Instrument Facility (CIF) of the Bose Institute and the Center for Cellular & Molecular Biology (CCMB), India.

## Author contributions

SM: conceptualization, formal analysis, investigation, visualization, methodology, and writing—original draft, review, and editing. DR: analysis and visualization. DB and Sriparna Mondal: investigation. KKS: biochemical assay. UB and SS: review. Anupama Ghosh: resources, supervision, writing—original draft, review, and editing. Abhrajyoti Ghosh: conceptualization, resources, supervision, funding acquisition, validation, project administration, and writing—original draft, review, and editing.

## Data availability

All data generated in this study were submitted to the NCBI (SRA) under the Bioproject numbers PRJNA1535431 and PRJNA1525924 and are freely available to the scientific community.

## Supplementary Figures

**Supplementary Fig. 1.**
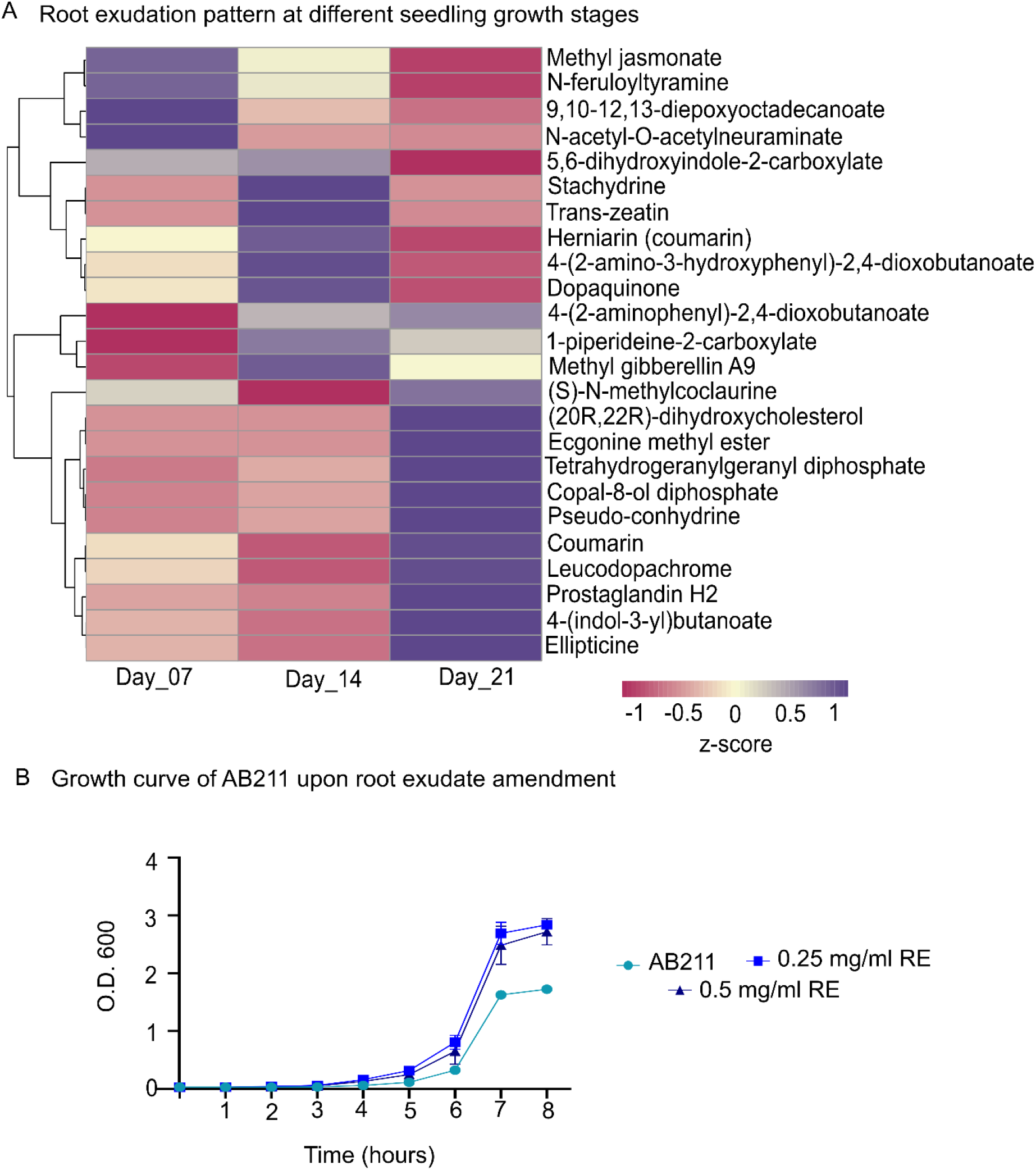
(A) Highly abundant metabolites in maize root exudates detected through LC MS/MS analysis (B) Planktonic growth of AB211 in the presence and absence of 0.25 and 0.5 mg/ml maize root exudate.

**Supplementary Fig. 2.**
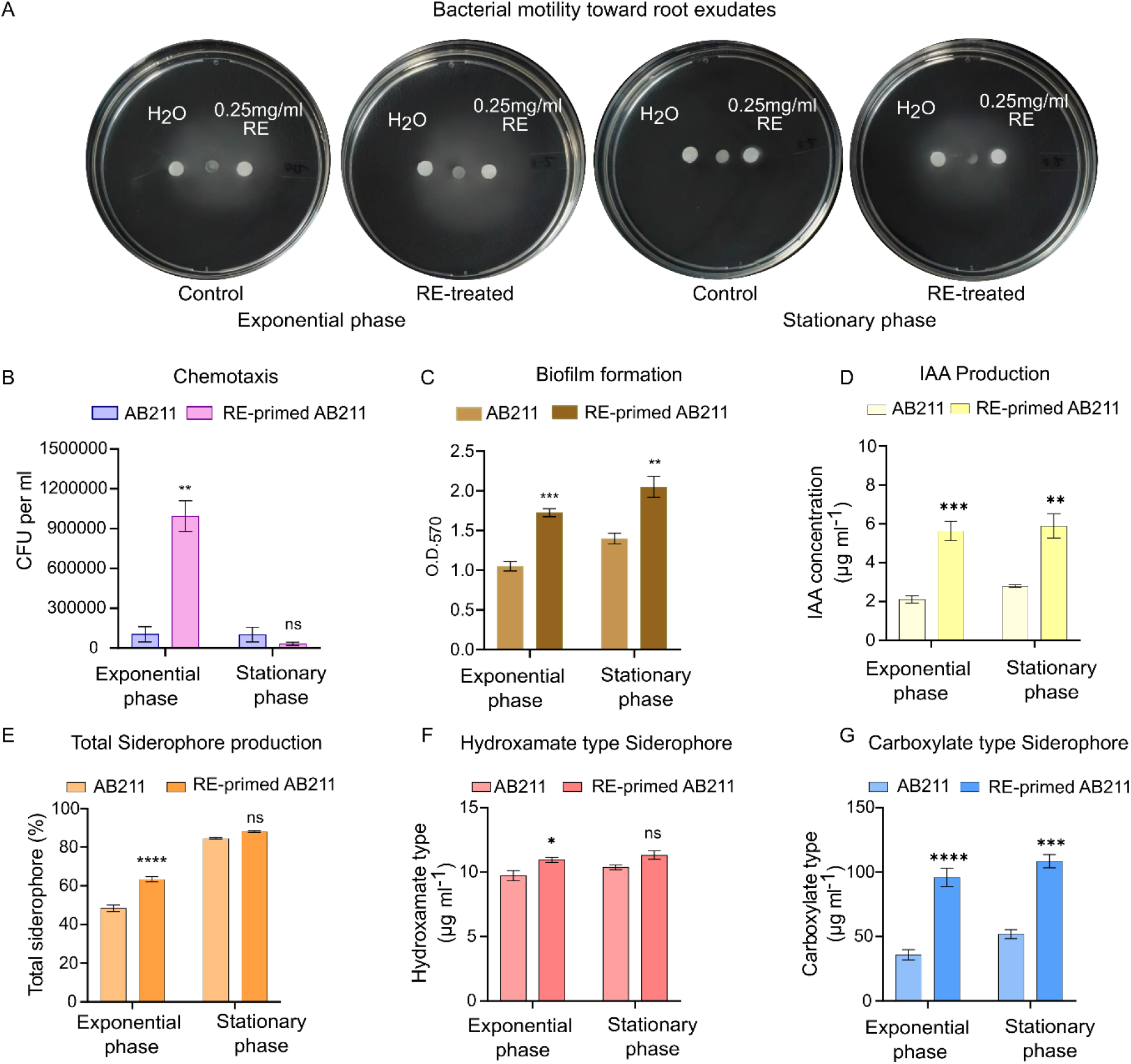
Physiological impact of root exudates on *P. aryabhattai* AB211. (A) Chemotaxis behavior of AB211 towards root exudates (0.25 mg/ml) during exponential phase and stationary phase. In both the cases, diffused movement of AB211 is more while treated with root exudates than the control cells. (B) Quantification of cells attracted towards root exudates using CFU count. (C) Biofilm formation, (D) IAA production, (E) total siderophore production, (F) Hydroxamate and (G) Carboxylate type siderophore production ability of AB211 upon root exudates exposure was biochemically quantified.

**Supplementary Fig. 3.**
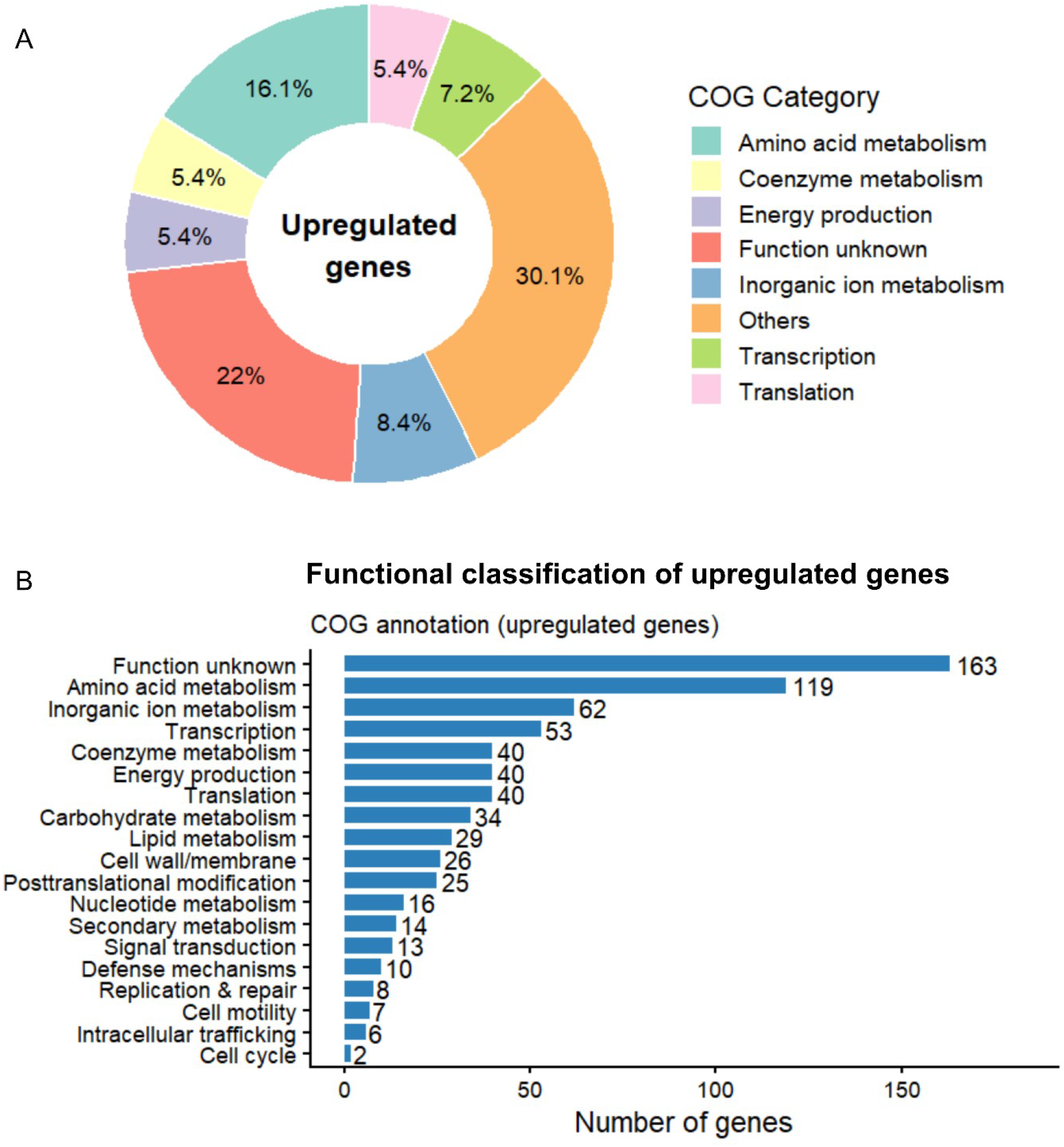
COG category disctribution. (A) and number of significantly upregulated genes under such categories (B) from transcriptome.

**Supplementary Fig. 4.**
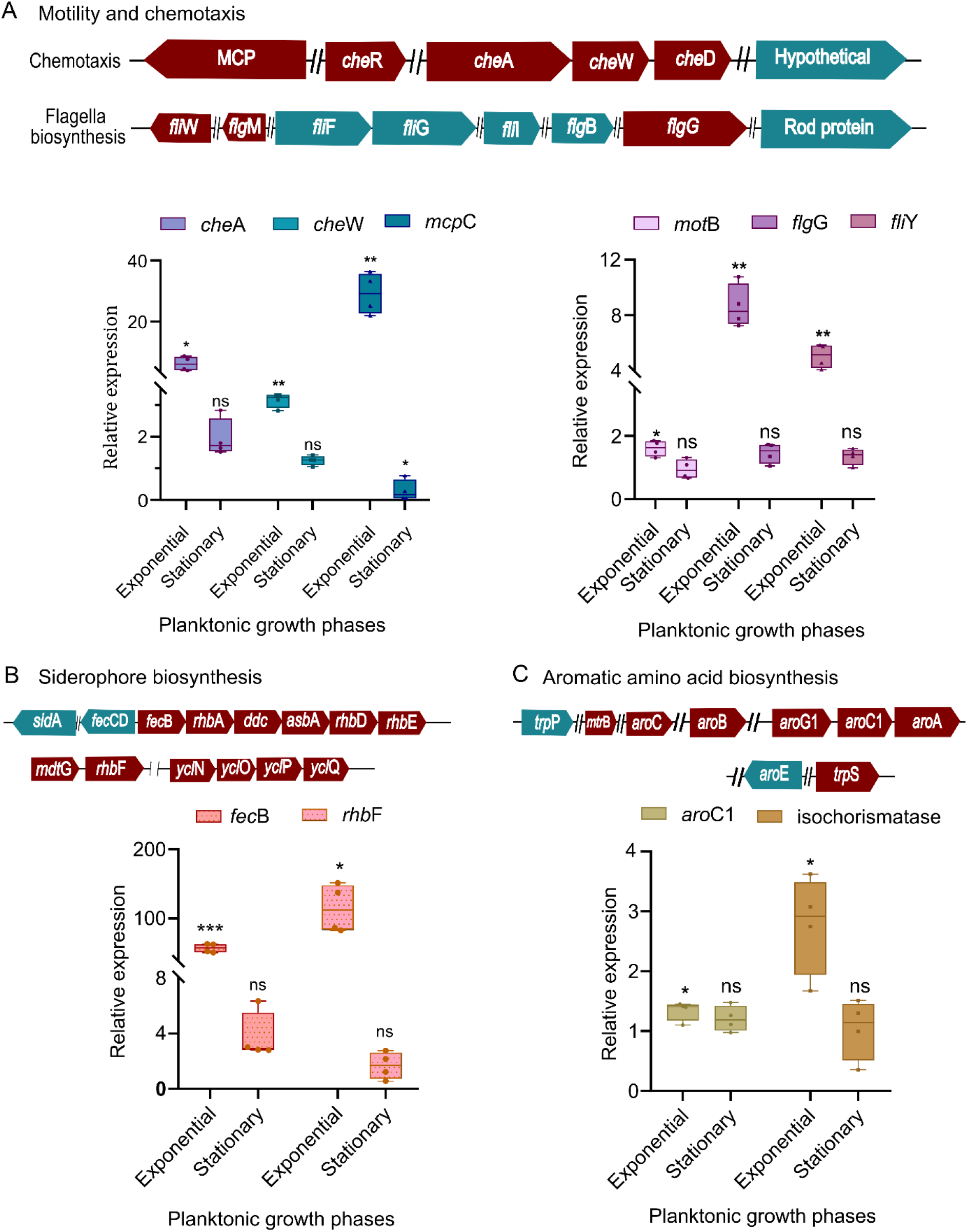
Real time expression of profile of genes involved in chemotaxis and motility. (A), siderophore (B) and aromatic amino acid biosynthesis (C) with the respective operon structure. One-way ANOVA was performed to evaluate the significance of the differences observed (pvalue ≤0.05).

**Supplementary Fig. 5.**
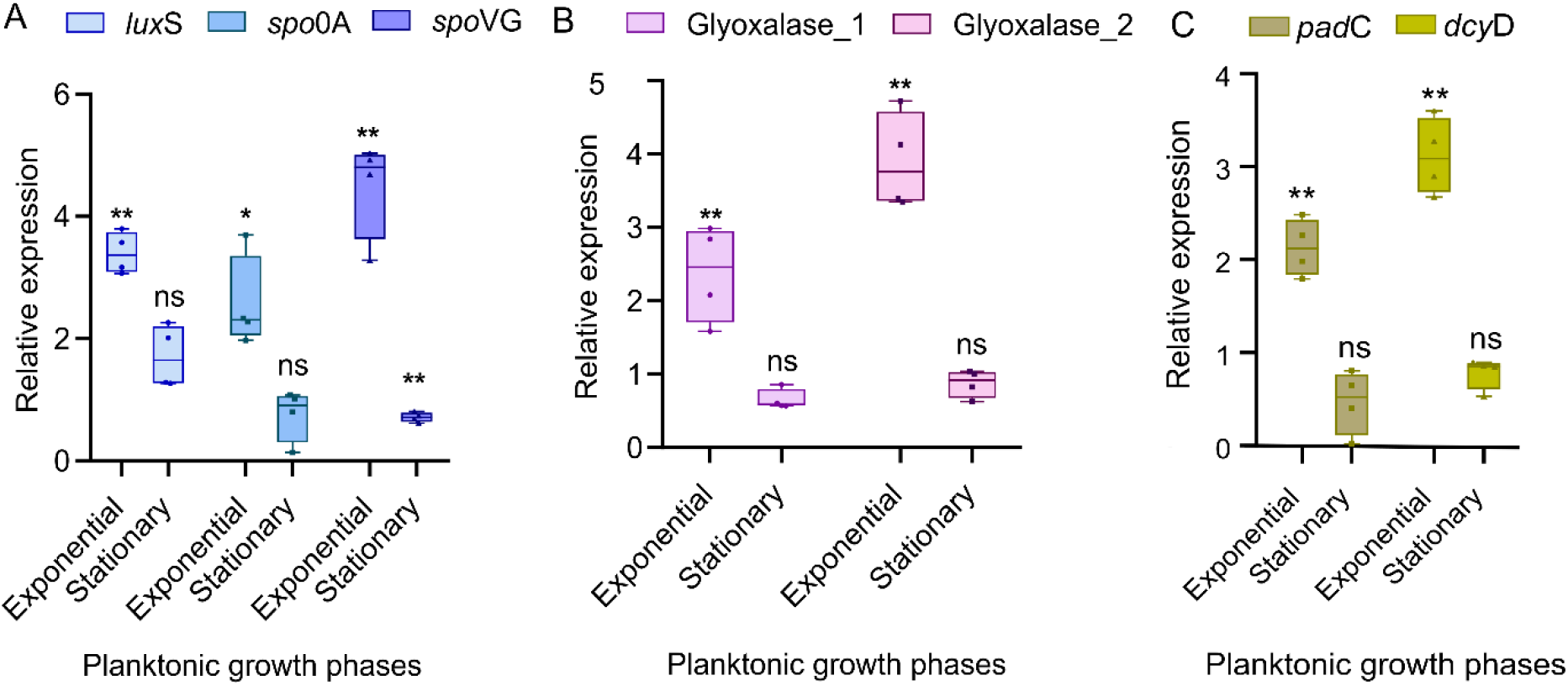
Real time expression of profile of genes involved in quorum sensing. (A), glyoxalate degradation (B) and phenolic acid associated pathways (C). One-way ANOVA was performed to evaluate their significance levels (pvalue ≤0.05).

**Supplementary Fig. 6.**
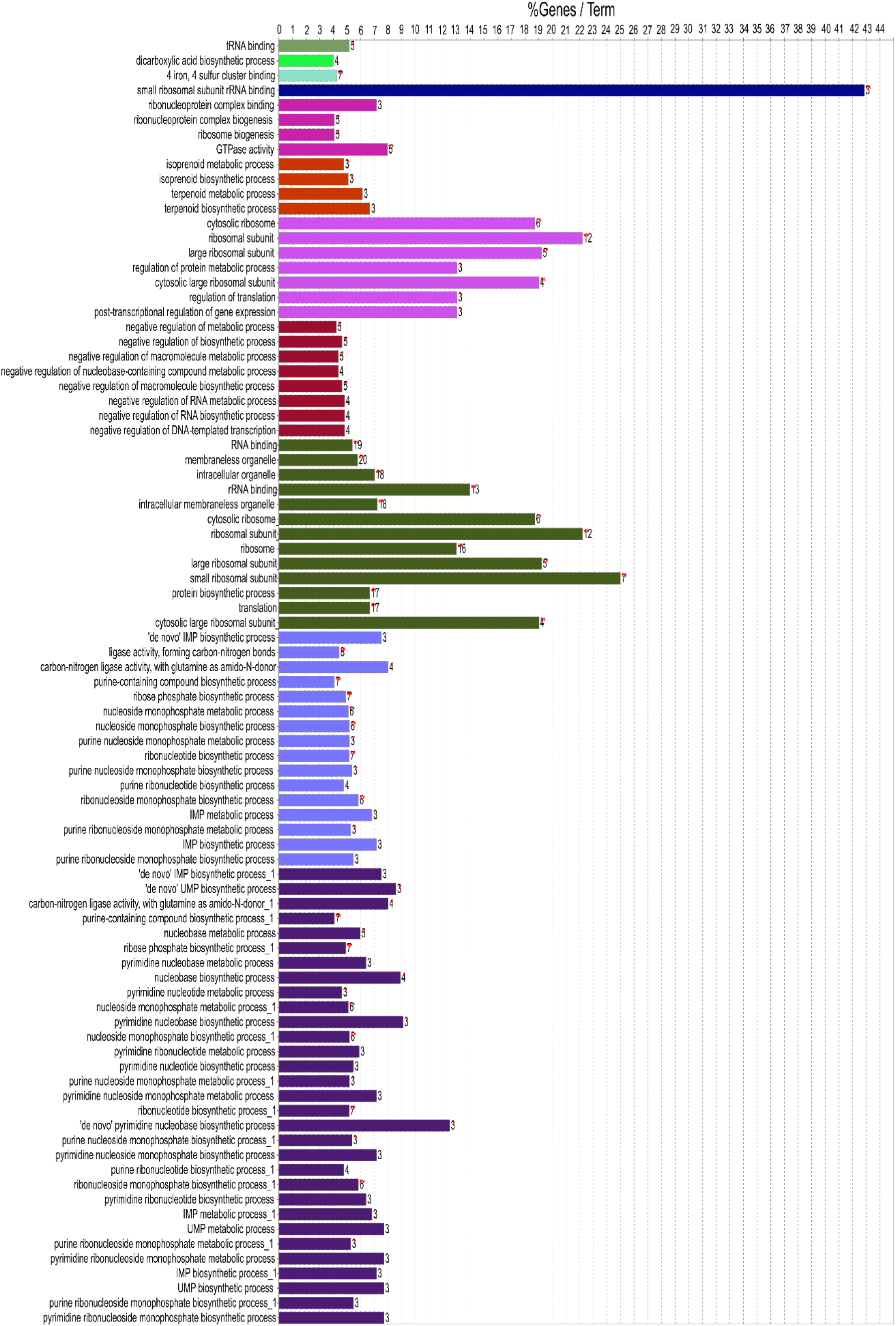
Functional enrichment of significantly altered proteins in AB211.

**Supplementary Fig. 7.**
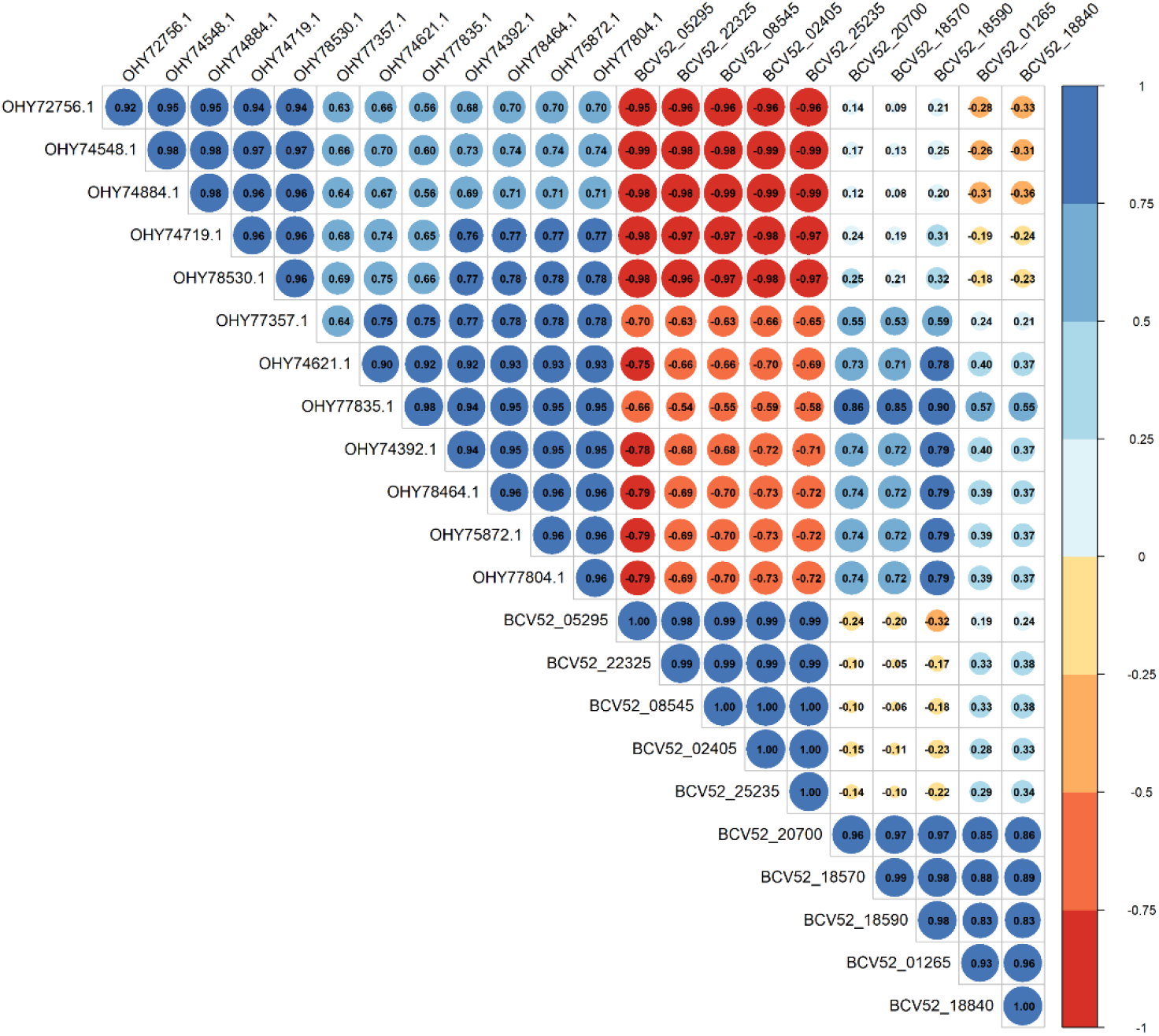
DIABLO-based multi-omics integration showing relationships between upregulated proteins (Log fold change ≥ 0.5, p-value ≤0.05) and transcriptionally regulated genes. Correlation matric showing positive (blue) and negative (red) correlation between the features, including the circle sizes reflecting correlation strength.

**Supplementary Fig. 8.**
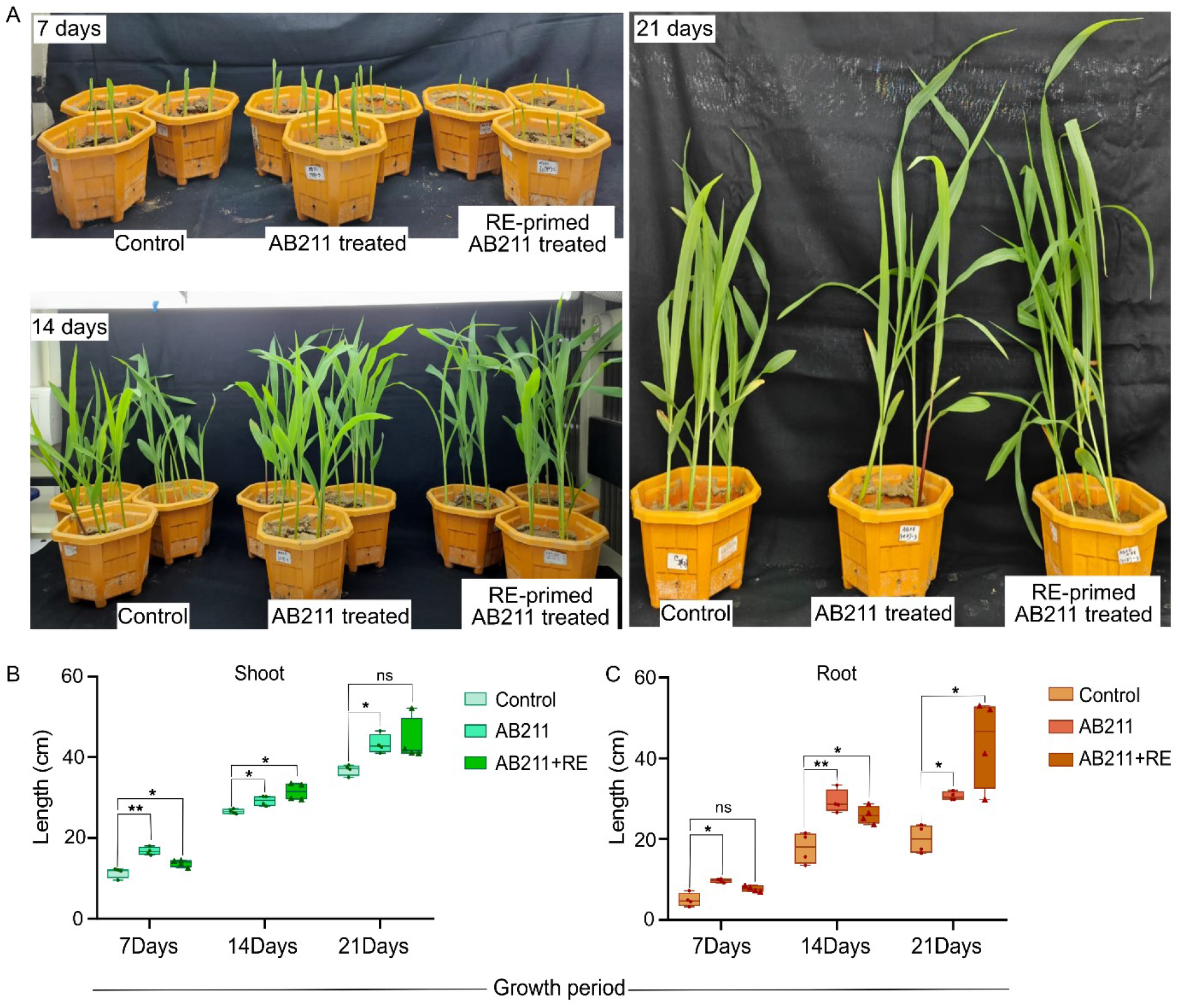
Plant growth-promotion upon AB211 amendment. Statistical significance of (B) shoot and (C) root length (cm) was calculated by two-way ANOVA (pvalue ≤0.05) followed by Tukey’s multiple comparisons test (n=4).

**Supplementary Fig. 9.**
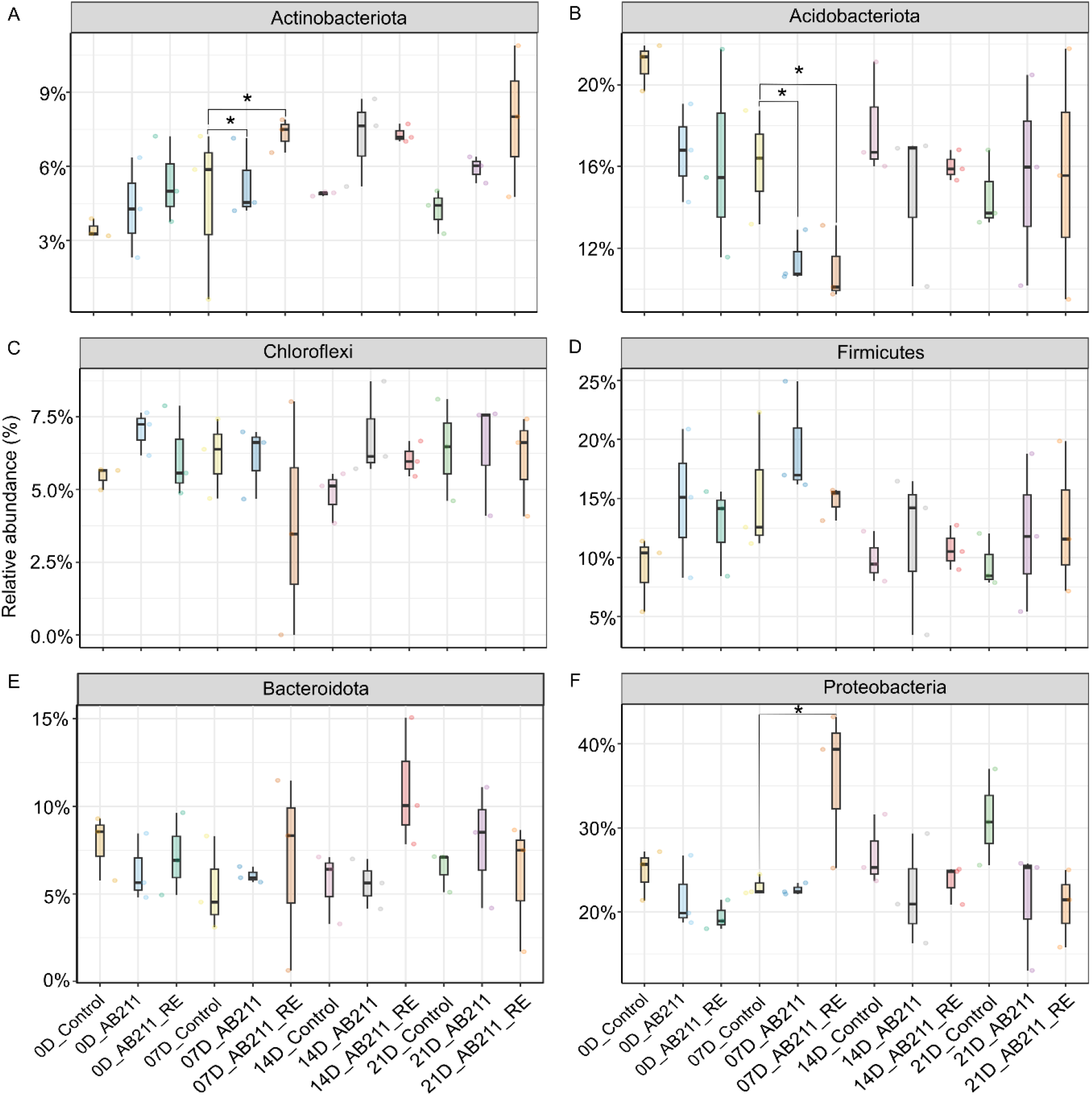
Distribution profile of actinobacteriota. (A), acidobacteriota (B), chloroflexi (C), firmicutes (D), bacteroidota (E) and proteobacteriota (F) throughout the experimental period. Pairwise t.test was performed to validate the significance of such a pattern, (*, p-value ≤ 0.05) and (ns, non-significant).

**Supplementary Fig. 10.**
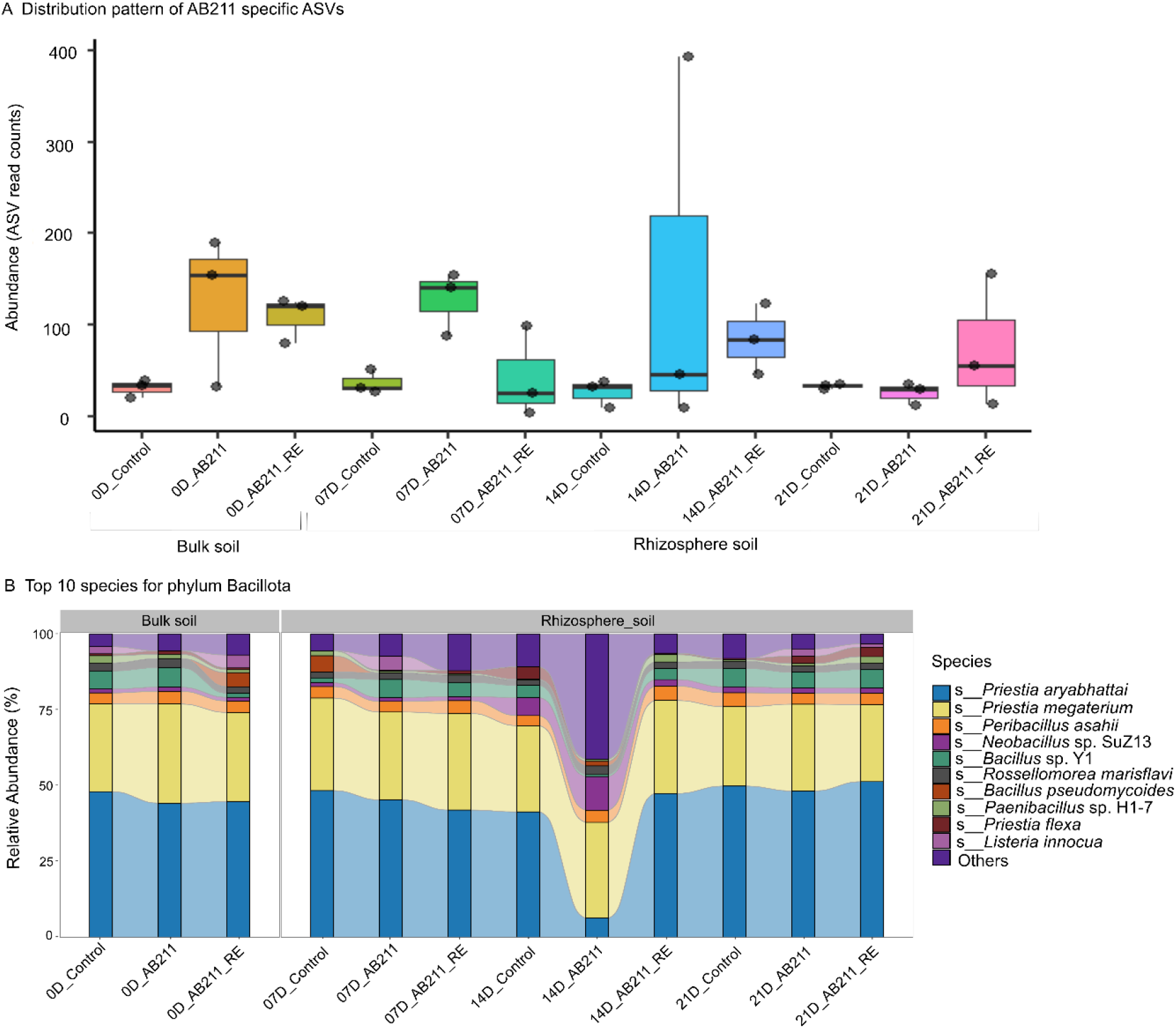
Distribution profile of AB211-associated OTUs (B) from amplicon sequencing and Bacillota group (C) from metagenomics.

